# Stoichiometric transcription factor partnerships control GABAergic neuron fate allocation

**DOI:** 10.64898/2026.05.25.727662

**Authors:** Elena Dvoretskova, Connor Lynch, Ann Rose Bright, Yigit Koray Babal, Sergey Isaev, Peter Kharchenko, Igor Adameyko, Christian Mayer

## Abstract

Combinatorial transcription factor (TF) codes specify neuronal fates, yet how quantitative differences in interacting TFs shape these decisions remains unresolved. We address this in the developing basal ganglia, where a pool of undifferentiated progenitors gives rise to both D1 and D2 medium spiny neurons (MSNs). Combining sparse *in vivo* CRISPR perturbation, lineage barcoding, and single-cell transcriptomics in mice, we find that loss of *Sp9* shifts clonal fate bias from D2 toward D1 MSNs and intercalated cells. Mechanistically, chromatin profiling and biochemical assays show that at GC-rich promoters, SP9 binds DNA directly and activates transcription. At distal enhancers, SP9 binds indirectly, tethered by DLX factors, represses activity, and associates with the NuRD corepressor complex. The relative abundance of SP9 and DLX selects between these modes. These findings extend the combinatorial TF code beyond factor identity to relative proportions, with relevance to neurodevelopmental disorders.

## Introduction

Most inhibitory neurons of the mammalian forebrain, which produce the neurotransmitter gamma-aminobutyric acid (GABA), originate from the embryonic ganglionic eminence (GE), a transient ventral telencephalic structure. The GE is subdivided into medial (MGE), lateral (LGE), and caudal (CGE) regions: the MGE and CGE produce most of the forebrain interneurons (INs), while the LGE produces the GABAergic projection neurons (PNs) of the basal ganglia and olfactory bulb (OB) neurons (Bandler and Mayer, 2023; van Velthoven et al., 2025; Flames et al., 2007). The principal striatal PNs are the medium spiny neurons (MSNs), which integrate cortical input and form the main striatal output through two subtypes defined by dopamine receptor expression: direct-pathway D1-type medium spiny neurons (D1 MSNs) and indirect-pathway D2-type medium spiny neurons (D2 MSNs) (Gerfen, 2022; Johansen et al., 2025). Recent integrative analyses identify the GE as a convergence point for the developmental origins of neurodevelopmental disorders (NDDs): many risk genes for autism spectrum disorder (ASD), intellectual disability, and epilepsy are enriched in GE progenitors and their immediate descendants (Aivazidis et al., 2025; Satterstrom et al., 2020; Yuan et al., 2025). These observations highlight the need to define how transcriptional regulatory mechanisms allocate GE-derived neuronal fates.

GABAergic diversity in the GE emerges through morphogen-mediated regional patterning that establishes region-specific transcriptional programs, with DLX homeobox transcription factors (TFs) functioning as master regulators of inhibitory neuron identity (Lindtner et al., 2019; Rubenstein et al., 2024). In mice, *Dlx* transcripts and DLX proteins are first expressed in GE progenitors at embryonic day (E)9.5 and follow a temporal sequence of DLX2, DLX1, DLX5, and subsequently DLX6 expression (Eisenstat et al., 1999; Rubenstein et al., 2024). DLX TFs regulate downstream gene expression by binding the TAATT motif within evolutionarily conserved regulatory elements (REs) (Catta-Preta et al., 2025; Lindtner et al., 2019). The TAATT motif, however, is broadly distributed across the genome, raising the question of how DLX-bound regulatory landscapes give rise to lineage-specific transcriptional outputs. One likely mechanism is combinatorial control through interactions with other TFs (Catta-Preta et al., 2025; Xie et al., 2025; Dvoretskova et al., 2024; Jolma et al., 2015). How such TF partnerships are organized at shared REs, and how quantitative differences in TF abundance are converted into subtype-specific fate allocation, remains unresolved.

To address this, we focused on the NDD-associated TF SP9, which belongs to the C2H2 zinc finger (ZF) family of Krüppel-like factors (KLF) and specificity proteins (SP) (Presnell et al., 2015). SP9 provides an entry point because it binds many of the same REs as DLX TFs in the GE (Catta-Preta et al., 2025), plays a central role in cortical interneuron development (Liu et al., 2019), and is required for the development of D2 MSNs (Zhang et al., 2016; Xu et al., 2018; Li et al., 2022), although the regulatory mechanism underlying this requirement has remained unresolved. In humans, *SP9* variants have recently been implicated in NDDs including epilepsy and intellectual disability (Tessarech et al., 2024). Because SP9 co-occupies REs with DLX TFs yet is required for a specific neuronal fate, it provides a tractable model to dissect how partner TFs allocate related GABAergic neuron fates.

Here, by combining sparse in utero electroporation (IUE)-based CRISPR-Cas9 perturbation in the embryonic GE with heritable single-cell lineage barcoding and continuous clonal-state embedding (Erickson et al., 2025; Isaev et al., 2026), we show that loss of *Sp9* redirects individual progenitors from D2 MSN fate toward D1 MSN and intercalated cell of the amygdala (ITC) fates. Mechanistically, we resolve SP9 activity into two distinct regulatory modes. At DLX-motif-containing distal REs, SP9 occupancy depends on its ZF-mediated interaction with DLX factors rather than on direct DNA contact; at these sites, SP9 associates with the nucleosome remodeling and deacetylase (NuRD) complex to repress enhancer activity. At a smaller set of GC-motif-rich promoters, including *Six3* and *Sp9* itself, SP9 binds directly and activates transcription through its N-terminal domain. The relative abundance of SP9 and DLX factors, rather than their individual levels, determines SP9 binding distribution between these two modes, with SP9 self-activation linking this regulatory balance to divergent D2 MSN versus D1 MSN fate allocation. Together, these results establish stoichiometric TF partnerships as a quantitative principle of developmental neuronal subtype specification.

## Results

### Sparse *in vivo* perturbation of *Sp9* redirects LGE projection neuron fate from D2 MSNs to D1 MSNs

To test whether *Sp9* specifies D2 MSN fate and to follow the cellular trajectories of perturbed progenitors, we combined sparse *in vivo* CRISPR perturbation (tCrop-seq) with heritable lineage barcoding (TrackerSeq) in the embryonic GE (Figure 1A). IUE at E12.5 broadly targeted progenitors across the medial, lateral, and caudal GE. We co-electroporated a single-guide RNA (gRNA) against *Sp9* (gSp9) or *LacZ* (gLacZ control; one gRNA per embryo), together with a barcode library that labels individual progenitors with random heritable barcode combinations, transducing a small fraction of cycling Cas9-expressing progenitors (Dvoretskova et al., 2024; Bandler et al., 2022). Sparse perturbation places gSp9 cells alongside unperturbed neighbors, enabling cell-autonomous effects to be assessed without confounds from altered tissue context. Daughter cells inheriting the same gRNA share a perturbation condition (gSp9 or gLacZ); daughter cells inheriting the same barcode constitute a clone. At E16.5, cortices, striata, and OBs were dissected, dissociated, and tdTomato^+^ cells were recovered by fluorescence-activated cell sorting (FACS) and profiled by single-cell RNA sequencing (scRNA-seq) to obtain transcriptomes, gRNA assignments, and clonal assignments at single-cell resolution.

**Figure 1:**
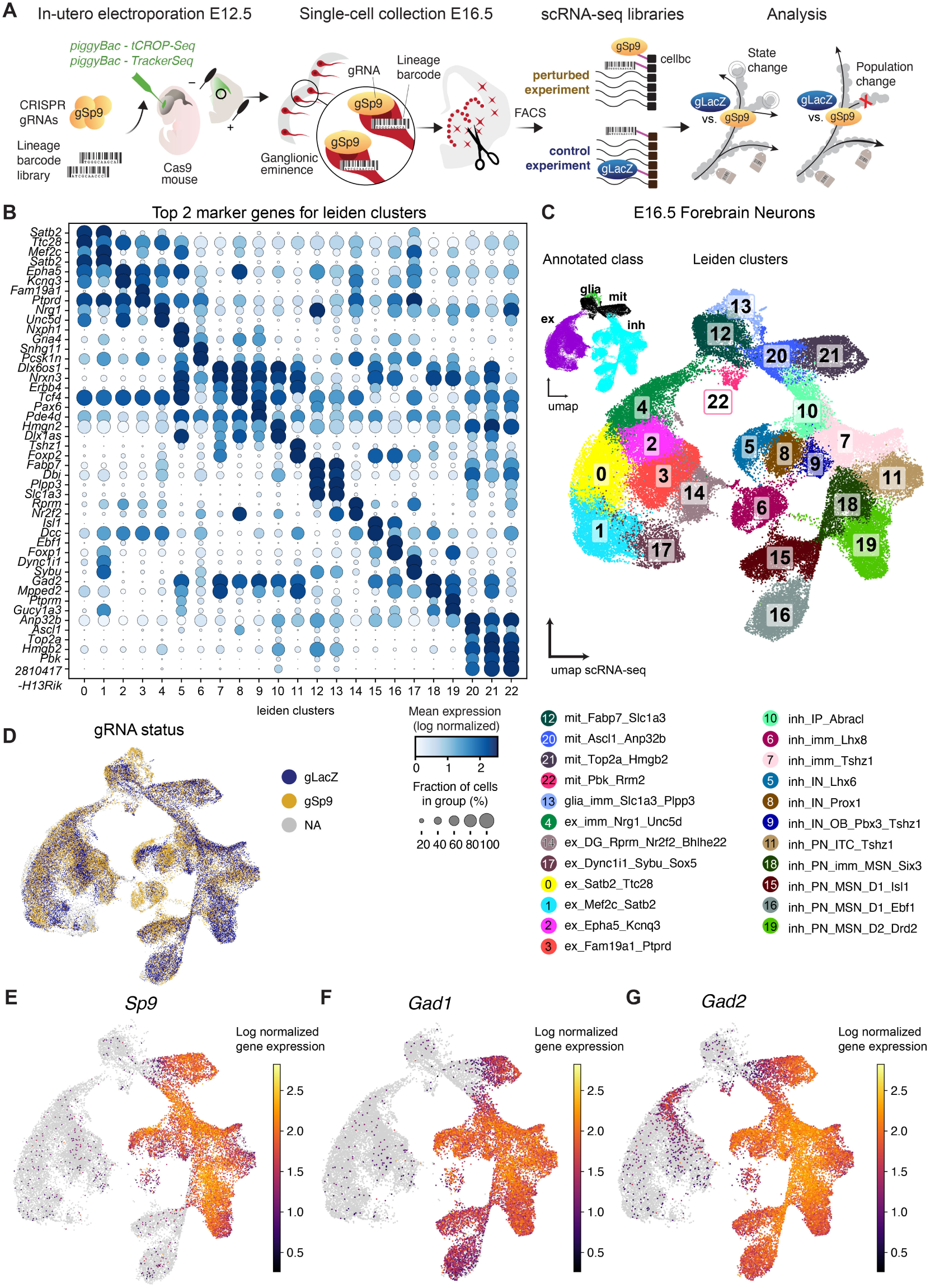
In vivo tCROP-seq perturbation of *Sp9* combined with TrackerSeq lineage tracing in the embryonic mouse ganglionic eminence. **A**, Design of the tCROP-seq sparse knockout plus TrackerSeq lineage tracing experiment. Either control (gLacZ) or perturbation (gSp9) guide RNAs (one type per embryo to prevent co-perturbation within individual clones), along with a diverse DNA barcode library were electroporated *in utero* into ganglionic eminences of embryonic day 12.5 (E12.5) Cas9-expressing embryos. Cells were collected at E16.5 and enriched by FACS for tdTomato (indicating gRNA expression). Single-cell RNA sequencing (scRNA-seq) jointly recovers whole transcriptomes, gRNA assignments, and lineage barcodes per cell, enabling analysis of how *Sp9* loss alters transcriptomic states and clonal lineage allocation during GABAergic differentiation. **B**, Top 2 marker genes per cluster from a single-cell Wilcoxon rank-sum test for Leiden clusters covering all recovered mitotic, glial, and neuronal cells. **C**, UMAP projections of mitotic, glial, and neuronal cells. Upper left: annotated classes (excitatory neurons, inhibitory neurons, mitotic cells, glia). Main panel: Leiden clusters at resolution 1.0 with manual cell-type annotations ordered by class, maturity status, subclass, cell type (where transcriptomically apparent), and top marker gene. **D**, UMAP showing the gRNA recovered per cell (gLacZ, gSp9, or NA). NA cells are those for which no gRNA transcript was detected, likely due to technical dropout during single-cell capture or sequencing. **E-G**, Feature plots of *Sp9* (E), the GABAergic fate marker *Gad1* (F), and the GABAergic fate marker *Gad2* (G). Counts are scaled and log-normalized.

After quality control and removal of vascular and microglia populations (Figure S1A-F), we recovered 73,754 cells (Table S1). Using Leiden clustering and canonical marker gene expression, we identified GABAergic neurons, mitotic progenitors, immature glia, and a transcriptionally distinct population of excitatory neurons from dorsal progenitors co-targeted at the cortex/GE border (Figure 1B-C). Across all recovered cells, 29,109 were assigned to gSp9 and 28,607 to gLacZ; the latter were combined with a published gLacZ dataset (Dvoretskova et al., 2024). The remaining cells had insufficient gRNA reads for confident assignment and were used only for cell-type annotation (Figure 1D). *Sp9* expression was restricted to the GABAergic lineage (Figure 1E-G), and subsequent perturbation analyses focused on GABAergic and mitotic cells. These cells partitioned into 28 transcriptomic states spanning cycling progenitors, postmitotic intermediates, and differentiating INs and PNs (Figures 2A, S2A-C); both gSp9 and gLacZ conditions were represented across all states (Figure 2B). Consistent with the known origin of D1 and D2 MSNs from a common progenitor pool, trajectory analysis (PAGA, diffusion pseudotime (Wolf et al., 2019)) recovered a bifurcating MSN lineage with shared progenitor states preceding D1 MSN and D2 MSN fate (Figure 2C-E) (Bandler et al., 2022).

**Figure 2:**
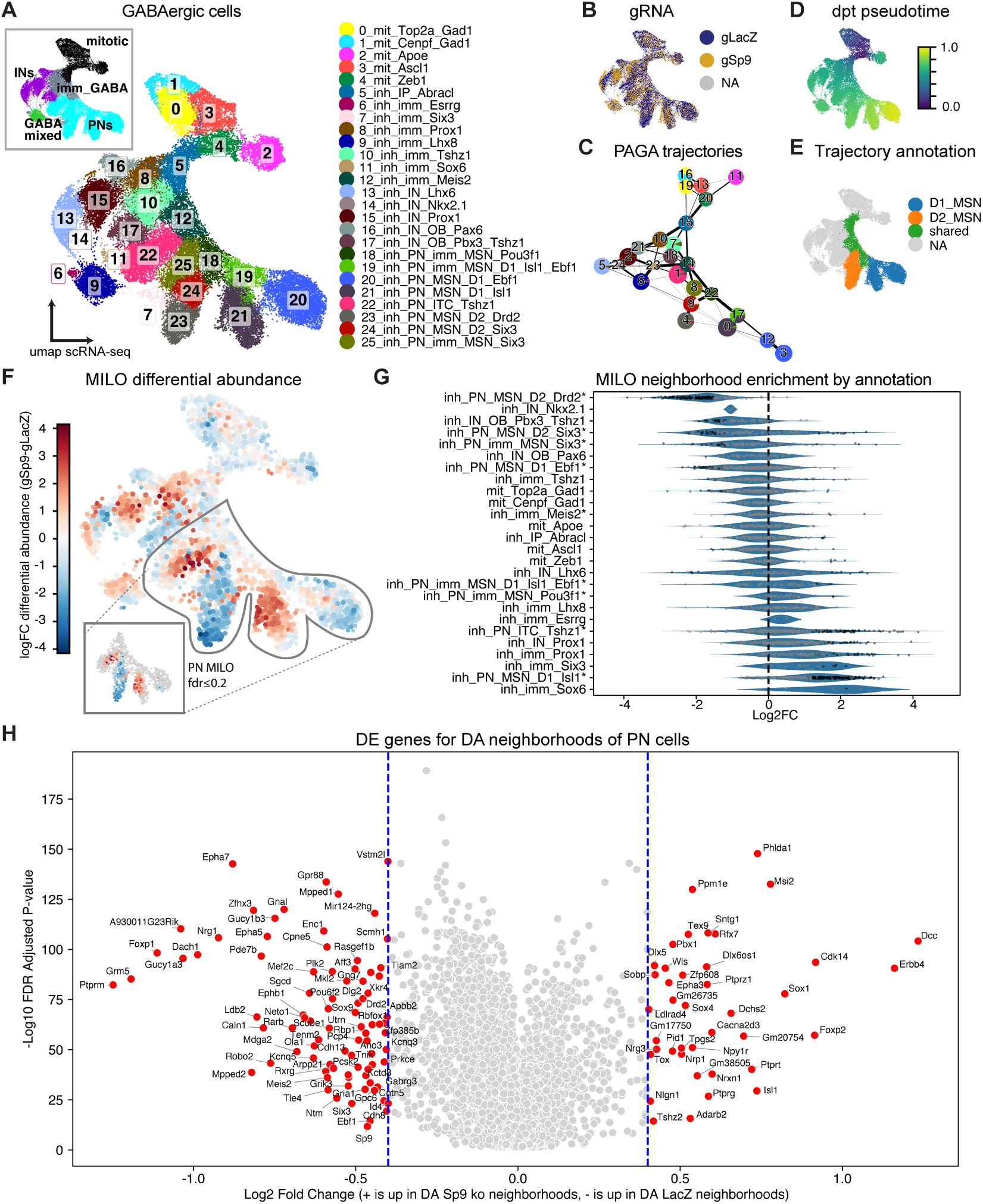
*Sp9* perturbation depletes D2 MSNs and enriches D1 MSNs and ITCs *in vivo*. **A**, UMAP projection of mitotic and GABAergic cells with cell subclass labels and Leiden cluster numbers at resolution 1.5. Cluster annotations were updated using refined markers as in Figure S2. **B**, UMAP showing the gRNA recovered per cell (gLacZ, gSp9, or NA). **C**, Partition-based graph abstraction (PAGA) trajectory estimation, showing GABAergic differentiation into interneuron (IN) and projection neuron (PN) branches, with PN precursors subsequently bifurcating into D1 and D2 medium spiny neuron (MSN) trajectories. Numbers in the PAGA graph denote Leiden cluster identities (matching the numbered clusters in Figure S2A). **D**, Geodesic diffusion pseudotime inference on the UMAP, with a cell from the inh_IP_Abracl cluster used as root node. **E**, Classification of cluster trajectories for D1 and D2 MSNs, with shared and distinct portions of the trajectory labeled based on the PAGA analysis in C. **F**, Differential abundance (DA) testing (MILO; knn = 100) across mitotic and GABAergic cells. Neighborhoods enriched for gSp9 cells compared to gLacZ cells are shown in red, depleted neighborhoods in blue. The outlined region contains the PN neighborhoods used for the differential expression analysis in panel H. Inset shows neighborhoods meeting a false discovery rate (FDR) threshold of ≤ 0.2. **G**, Beeswarm plot of MILO neighborhood log_2_ fold-change in abundance (gSp9 vs gLacZ), grouped by majority cell-type identity. Neighborhoods shown in black meet the FDR ≤ 0.2 threshold. Categories marked with an asterisk denote neighborhoods used in the PN neighborhood differential expression analysis. **H**, Volcano plot of differentially expressed (DE) genes between significantly enriched versus depleted PN neighborhoods. Genes meeting | log_2_ FC| ≥ 0.4 and FDR ≤ 0.2 are highlighted in red.

We focus on the LGE-derived projection neuron lineage. Within each replicate, we compared gSp9 with gLacZ to isolate perturbation-specific effects. IN-related changes are reported in supplementary figures. Differential abundance testing on the GABAergic dataset using MILO (Dann et al., 2022) revealed the strongest changes within the LGE neuron lineage: gSp9 neighborhoods were significantly depleted of D2 MSNs and enriched for *Isl1*^+^ D1 MSNs and *Tshz1*^+^ ITCs (Figure 2F,G). Changes in IN populations were also observed, with depletion of *Pbx3*^+^ OB INs and enrichment of *Prox1*^+^ cortical INs and *Sox6*^+^ immature INs (Figure S3A-B), consistent with the ectopic accumulation of *Sox6*^+^ cells in the ventral telencephalon previously reported in conditional *Sp9* knockouts (Liu et al., 2019).

The transcriptional signature of perturbed cells matched these abundance shifts. MILO neighborhood differential expression (DE) analysis on the PN-classified subset showed that gSp9-enriched neighborhoods upregulated D1 MSN and ITC markers (*Isl1*, *Foxp2*, *Sox6*, *Zeb2*, *Arx*, *Fign*, *Dlx6os1*, *Erbb4*) and downregulated D2 MSN markers (*Drd2*, *Penk*, *Adora2a*, *Gpr6*, *Six3*, *Meis2*, *Grik3*) (Figure 2H, Table S3). DE analysis of IN-containing neighborhoods identified additional changes in cell-guidance and synaptic genes, including several associated with epilepsy (*Gria1*, *Kcnd2*, *Nrxn1*, *Cacna1c*; Figure S3B-C (Piacentini et al., 2026)). Independent sample-pseudobulked DE analysis, comparing matched cell types across replicates, converged on the same directional signature and additionally identified genes not captured by neighborhood-level testing (*Ppm1j*, *Ptprt*, *Npy1r*, *Nrp1*) (Table S4, Figure S3D-E).

Together, the concordance of cell-abundance and gene-expression shifts indicates that *Sp9* loss reshapes the output of the LGE projection lineage.

### *Sp9* loss does not eliminate, but rather redirects, the LGE projection lineage

Population-level analyses cannot distinguish fate redirection from alternative mechanisms such as selective loss of prospective D2 MSNs or developmental arrest at a transitional state. Clonal lineage tracing (TrackerSeq) resolves this by linking each barcode-labeled progenitor to the set of cell types it produces. After filtering of clonal signals using Jaccard similarity and clone assignment, the dataset comprised 13,598 cells distributed across 5,485 multi-cell clones, with clone sizes ranging from 2 to 11 cells (Figures 3A, S4A-B).

**Figure 3:**
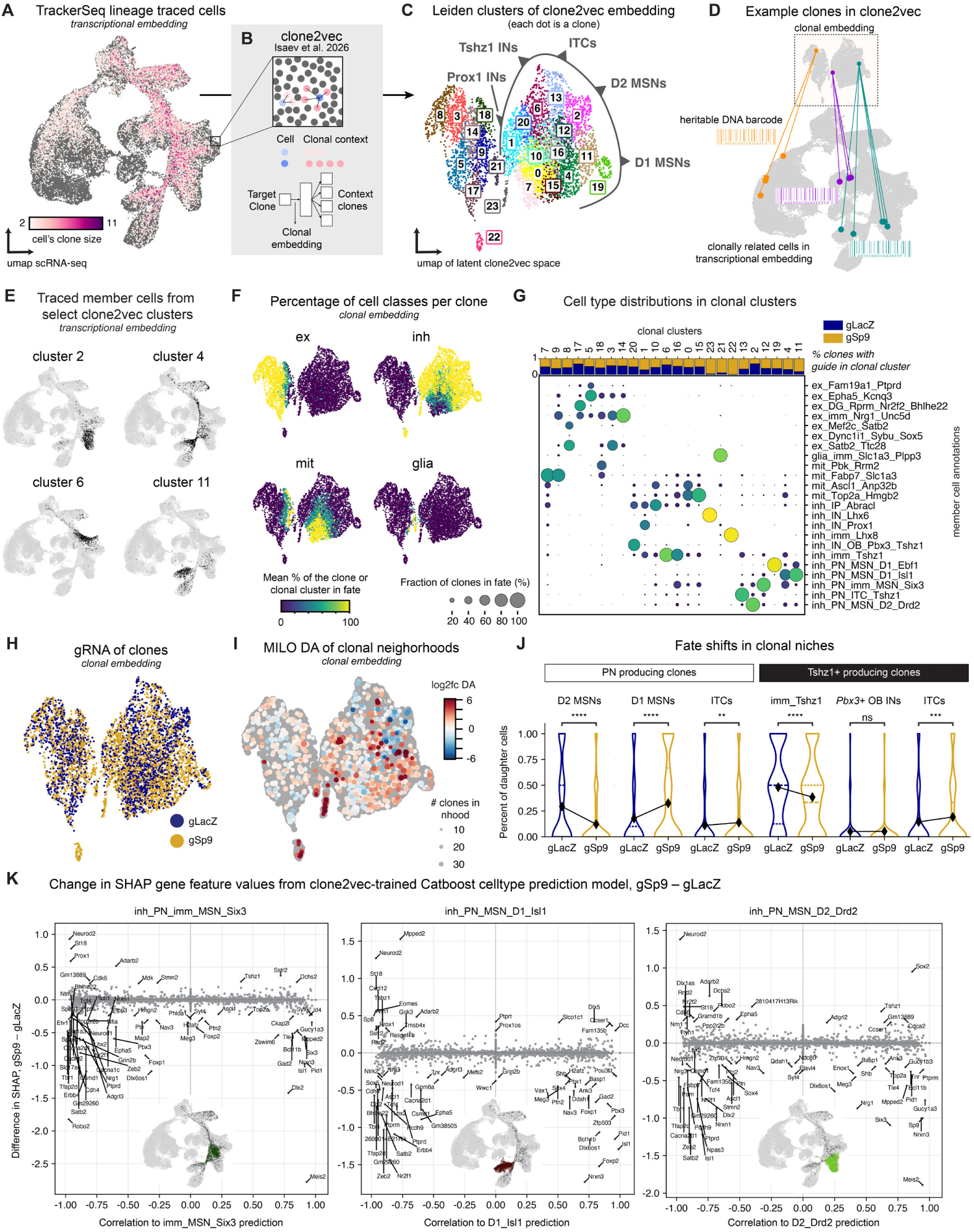
Clonal lineage tracing reveals *Sp9*-dependent fate biases in D1 and D2 MSN-producing clones. **A**, UMAP showing all cells assigned to multi-cell TrackerSeq clones, colored by the size of the clone each cell belongs to. **B**, Schematic of the clone2vec latent embedding of TrackerSeq clones, adapted from (Isaev et al., 2026). The cellular composition of input clones is used to predict context clones with similar transcriptomic compositions; the resulting vectorized clone embeddings provide a latent representation of the clonal dataset. **C**, Clone2vec latent embedding of all multi-cell clones from the gSp9 and gLacZ TrackerSeq dataset. Colors indicate Leiden clusters at resolution 1.5. Manual annotations indicate dominant clonal fate biases across the embedding. **D**, Three example clones shown in both clone2vec latent space (top) and scRNA-seq gene expression space (bottom). Heritable DNA barcodes link the clonal embedding to the corresponding cells in transcriptomic space. **E**, Member cells of clones from selected clone2vec Leiden clusters (2, 4, 6, 11), shown on the scRNA-seq UMAP. **F**, UMAPs of the clone2vec embedding, with each clone (dot) colored by the mean percentage of its member cells classified into one of four cell classes (excitatory, inhibitory, mitotic, glia). Many clones produce mixtures of classes, so percentages per class typically sum to less than 100%. **G**, Top: Proportion of clones per clone2vec cluster (x-axis) assigned to gLacZ or gSp9 conditions. Bottom: Dot plot showing the cell-type composition of clones in each clone2vec cluster (x-axis), with transcriptomic cell-type annotations from the scRNA-seq analysis (Figure 1) on the y-axis. Dot size denotes the fraction of clones in each clone2vec cluster producing the corresponding cell-type fate. **H**, Clone2vec embedding with clones colored by gRNA identity (gLacZ or gSp9). **I**, Differential abundance testing (MILO) on the clone2vec embedding, showing enrichment of clones producing D1 MSNs, ITCs, and Prox1+ INs in the gSp9 condition, and depletion of clones producing D2 MSNs and Tshz1+ OB INs. **J**, Cell-type composition shifts in clones producing at least one PN cell (left) or at least one Tshz1+ cell (right), upon *Sp9* perturbation. *P*-values from Brunner-Munzel test; not significant (ns), p > 0.05; ^∗^p ≤ 0.05; ^∗∗^p ≤ 0.01; ^∗∗∗^p ≤ 0.001; ^∗∗∗∗^p ≤ 0.0001. **K**, Differences in gene SHAP values from a CatBoost cell-type prediction model trained on the clone2vec embedding, comparing gSp9 vs gLacZ conditions for three target cell types: inh_PN_imm_MSN_Six3 (left), inh_PN_MSN_D1_Isl1 (middle), and inh_PN_MSN_D2_Drd2 (right). The x-axis shows the correlation of each gene with the corresponding cell-type prediction in the gLacZ model; the y-axis shows the change in SHAP value (gSp9 minus gLacZ). Insets show on the scRNA-seq UMAP the cells belonging to clones that produced cells in the highlighted target cluster.

To analyze patterns of clonal fate variation, we applied clone2vec (Erickson et al., 2025; Isaev et al., 2026), which embeds each clone as a vector in a latent space based on the transcriptomic states of its member cells (Figure 3B). This representation captures continuous variation in clonal fate biases. Visualization of the embedding by uniform manifold approximation and projection (UMAP) and unbiased clustering identified groups of clones with similar cell-type composition (Figures 3C, S4C). Since the cells of a clone may span developmental stages and divergent differentiation outcomes, member cells of a single clonal cluster typically occupied multiple transcriptomic clusters (Figure 3D-E); the clonal embedding thus groups clones according to the composition of cell types they produce rather than the transcriptomic state of any individual cell.

At the broadest level, clones segregated into discrete niches corresponding to major lineage classes — inhibitory GABAergic, *Lhx8*^+^ cholinergic, proglial, and excitatory glutamatergic — together with their associated mitotic progenitors (Figure 3F). Within each niche, sub-lineages were arranged along continuous axes of variation, reflecting the graded nature of fate decisions in the forebrain (Figure 3C). Quantifying the cell types produced by each clonal cluster confirmed this pattern (Figure 3G): some clusters showed largely unimodal fate output (*Lhx6*^+^ INs, *Ebf1*^+^ D1 MSNs), while clusters producing *Isl1*^+^ D1 MSNs, D2 MSNs, ITCs, and *Prox1*^+^ or *Tshz1*^+^ INs exhibited graded biases with majority output of one type and minority production of related fates, consistent with multipotent rather than fate-restricted progenitors. Example clones plotted on both the clonal and transcriptomic embeddings illustrate this dual resolution (Figure 3D): sister cells from individual progenitors largely belonged to the same major lineage but varied along sub-lineage axes such as D1 MSN versus D2 MSN bias.

Differential abundance testing on the clonal embedding showed depletion of D2 MSN-producing clones in the gSp9 condition, accompanied by enrichment of clones biased toward *Isl1*^+^ D1 MSNs and *Tshz1*^+^ ITCs (Figure 3H-I). Excitatory clones — derived from progenitors that do not express *Sp9* — showed little change in abundance, providing an additional internal negative control.

To test whether perturbed progenitors redirect their fate within the PN lineage, we restricted analysis to the 1,828 clones that produced at least one PN cell (D1 MSN, D2 MSN, or ITC) and quantified the proportion of each fate per clone. In gSp9 clones, the share of D2 MSN cells produced per clone decreased significantly, with a comparable increase in D1 MSN cells and a minimal increase in ITCs (Figure 3J). Clone sizes were comparable between gSp9 and gLacZ conditions (Figure S4B), arguing against differential survival as the primary driver of this shift. Together with the matched fate-marker expression changes (Figure 2H), these intra-clone compositional shifts are most consistent with fate redirection, in which the D2 MSN output normally generated within the LGE projection lineage is reallocated toward D1 MSN and, to a lesser extent, ITC fates upon *Sp9* loss. A complementary analysis restricted to clones producing at least one cell from the *Tshz1*^+^ clonal niche (immature *Tshz1*^+^ cells, ITCs, and *Pbx3*^+^ OB INs) showed that the ITC increase under gSp9 was partially mirrored by a reduction in immature *Tshz1*^+^ cells, while *Pbx3*^+^ OB INs were unchanged (Figure 3J).

Cells within a clone span a range of differentiation states, so gene expression in less-differentiated cells can be linked to the cell-type composition that the clone ultimately produces (Figure S4D-E). We trained a gradient-boosting model (CatBoost) on aggregated clonal gene expression to identify molecular features predictive of clonal fate (Erickson et al., 2025; Isaev et al., 2026). Intuitively, a gene whose expression best predicts the cell types a clone ultimately produces is a top predictive feature, and a gene that loses predictive weight when *Sp9* is absent is one whose link to fate depends on SP9. In control (gLacZ) clones, *Sp9*, *Dlx2*, *Meis2*, *Foxp1*, and *Foxp2* were the top predictive features for global GABAergic fate as well as D1 MSN and D2 MSN identity (Figure S5A). Comparison of CatBoost models trained separately on gLacZ versus gSp9 clones revealed that *Dlx2*, *Foxp2*, *Meis2*, and *Six3* (the latter a known master regulator of D2 MSN fate (Song et al., 2021)) broadly lost predictive weight across the shared MSN precursor state and the terminally fated D1 MSN and D2 MSN states (Figure 3K), with similar losses across additional GE-derived populations (Figure S5B). This pervasive dependence of fate-predictive features on *Sp9* suggests that SP9 acts within a combinatorial network of regulators including DLX TFs, motivating the molecular analyses below of how SP9 and DLX interact at shared REs.

### SP9 and DLX factors co-occupy distal REs, where their combined presence represses regulatory activity

To characterize the genomic binding of SP9, we analyzed published SP9 chromatin immunoprecipitation sequencing (ChIP-seq) data from the E13.5 mouse GE (Xu et al., 2018; Catta-Preta et al., 2025), identifying 4,032 SP9-bound regions. Peaks proximal to transcription start sites (TSSs) were annotated as promoter-like, whereas distal peaks overlapping published H3K4me1 and H3K27ac profiles (Lindtner et al., 2019) were annotated as enhancer-like. The majority (∼70%) of SP9 peaks were enhancer-like, while only ∼6% were promoter-like; the remainder did not meet criteria for either category (“Other”; Figure 4A).

**Figure 4:**
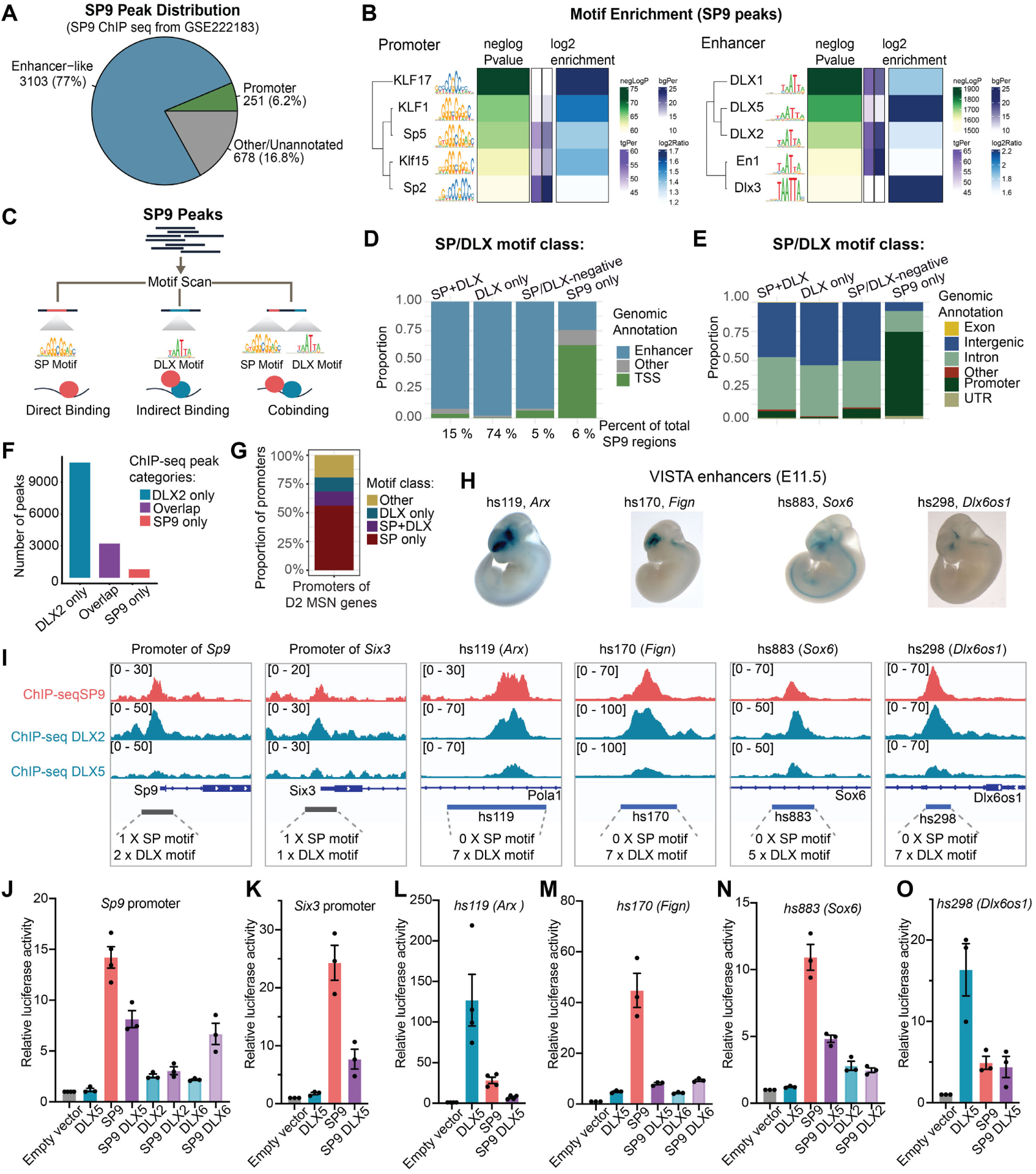
SP9 occupies DLX-motif-containing enhancers of GABAergic lineage genes through interaction with DLX factors. **A**, Genomic distribution of SP9 ChIP-seq peaks based on data from (Xu et al., 2018; Catta-Preta et al., 2025). Peaks localize predominantly to enhancer-like regions (77%), with smaller fractions at promoters (6.2%) and other regions (16.8%). **B**, Motif enrichment within SP9 peaks. Promoter-associated peaks (left) are enriched for GC-rich motifs recognized by KLF and SP factors, whereas enhancer-like peaks (right) are enriched for TAATT motifs recognized by DLX homeobox factors. Within each block: motif logos (left), − log_10_ *P*-value and target/background percentage of peaks containing the motif (middle), and log_2_ enrichment (right). **C**, Schematic of the motif scan-based classification of SP9 peaks. Peaks were grouped based on the presence of an SP motif only (direct binding), a DLX motif only (indirect binding via DLX), or both (cobinding). **D**, Coarse genomic annotation (enhancer, transcription start site (TSS), or other) of the four SP9 peak categories defined in (C): SP9+DLX (both motifs present), DLX (DLX motif only), Other, and SP9 (SP motif only). **E**, Fine genomic annotation (exon, intron, intergenic, promoter, UTR, other) of the same four peak categories. **F**, Intersection of DLX2 and SP9 ChIP-seq peaks from E13.5 mouse GE, based on data from (Lindtner et al., 2019; Catta-Preta et al., 2025; Xu et al., 2018). Bar plot shows the number of overlapping and non-overlapping peaks per category (DLX2 only, overlap, SP9 only). **G,** Motif class composition of promoters of D2 MSN marker genes that are downregulated upon *Sp9* perturbation (MILO neighborhoods, FDR ≤0.2) and carry an annotated SP9 ChIP-seq peak (n = 29 genes). SP-only and SP+DLX classes together account for the majority of promoters, consistent with direct SP9 binding at multiple D2 MSN fate genes beyond *Six3*. **H**,Whole-mount X-Gal staining (blue) of E11.5 transgenic mouse embryos carrying VISTA enhancer reporters hs119 (near *Arx*), hs170 (near *Fign*), hs883 (near *Sox6*), and hs298 (near *Dlx6os1*) used in the luciferase assays in panels L-O and shown as genome browser tracks in panel I. Images from (Visel et al., 2007). **I**, Genome browser tracks showing ChIP-seq signal for SP9, DLX2, and DLX5 in E13.5 mouse GE at the *Sp9* and *Six3* promoters (highlighted in grey) and the VISTA enhancers hs119, hs170, hs883, and hs298 (highlighted in blue). Numbers below each track indicate the count of SP and DLX motifs within the selected regulatory element. Data from (Lindtner et al., 2019; Catta-Preta et al., 2025; Xu et al., 2018). **J-O**, Luciferase reporter activity in N2A cells transfected with combinations of *Sp9* and *Dlx2*, *Dlx5*, or *Dlx6* as indicated, driven by the *Sp9* promoter (**J**), *Six3* promoter (**K**), or enhancer regions hs119 of *Arx* (**L**), hs170 of *Fign* (**M**), hs883 of *Sox6* (**N**), and hs298 of *Dlx6os1* (**O**). Bars represent mean ± s.e.m. of 9 (J,K,M-O) or 12 (L) replicates from 3 (J,K,M-O) or 4 (L) independent batches performed in triplicate; points indicate batch means. Statistical significance was assessed by two-way ANOVA with Tukey’s honestly significant difference (HSD) post hoc test. Exact *P*-values are provided in Table S7.

Motif enrichment showed that promoter-like SP9 peaks were enriched for GC-rich SP/KLF motifs, whereas enhancer-like peaks were enriched for the TAATT motif recognized by DLX TFs (Figure 4B), suggesting DLX-mediated indirect binding of SP9, which is consistent with prior reports of physical interactions between the SP and DLX TF families (Hojo et al., 2016; Castilla-Ibeas et al., 2025). Based on the presence of SP and DLX motifs, SP9-bound regions were classified into four classes (Figure 4C): DLX-only (DLX motifs but no SP motifs), SP-only (SP motifs but no DLX motifs), SP+DLX (both), and SP/DLX-negative (neither). Within the SP/DLX-negative class, the most enriched motif was the TGTCA motif recognized by MEIS1/MEIS2 (Table S5).

The motif classes mapped to distinct genomic features (Figure 4D,E): SP-only peaks were concentrated at TSSs and promoter regions (6 %), whereas DLX motif-containing peaks (DLX-only and SP+DLX; 74 % and 15 %) and SP/DLX-negative (5 %) peaks were enriched in enhancer, intronic, and intergenic regions. Comparison with DLX2 ChIP-seq peaks (from E13.5 GE; (Lindtner et al., 2019; Catta-Preta et al., 2025)) confirmed extensive co-occupancy: 3,226 of 4,032 SP9-bound regions were co-occupied by DLX2 (Figure 4F).

To examine the motif composition at SP9-bound promoters of D2 fate genes affected by *Sp9* perturbation, we focused on the top 100 D2 MSN markers downregulated upon *Sp9* loss. Of these, the 29 promoters with the highest SP9 ChIP-seq signal were subjected to motif analysis: the majority contained SP-family motifs, predominantly SP-only with a smaller subset also carrying DLX motifs (SP+DLX; Figure 4G), consistent with direct SP9 binding through its own SP motif at the promoters of many D2 MSN fate genes.

To probe the regulatory consequences of SP9 binding, we selected six REs for detailed analysis: two promoters of D2 MSN-associated genes (*Sp9* itself and *Six3*, a known master regulator of D2 MSN fate (Song et al., 2021)) and four VISTA enhancers (Visel et al., 2007) near D1 MSN marker genes upregulated in gSp9 MILO neighborhoods (hs119 near *Arx*, hs170 near *Fign*, hs883 near *Sox6*, and hs298 near *Dlx6os1*; Table S3), all of which drive enhancer activity in the embryonic forebrain at E11.5 (Figure 4H). All six REs showed co-binding of SP9, DLX2, and DLX5, but their motif content differed sharply: the *Sp9* and *Six3* promoters each contained a single SP motif alongside one or two DLX motifs, whereas all four enhancers contained multiple DLX motifs (5 to 7) but no SP motifs (Figure 4I). This asymmetry, with SP motifs present at the two promoters but not at the four distal enhancers tested, supports two distinct modes of SP9 engagement: direct DNA binding at promoters and motif-independent recruitment at enhancers.

To test the functional consequences of these binding modes, we performed dual-luciferase reporter assays in Neuro-2a (N2A) cells. SP9 alone activated reporter activity at the *Sp9* and *Six3* promoters and at several enhancers, and individual DLX factors (DLX2, DLX5, DLX6) similarly activated reporter activity at most REs. By contrast, co-expression of SP9 with any DLX factor significantly reduced reporter activity at all six REs relative to SP9 or the respective DLX factor alone (Figure 4J-O; Table S7). These results indicate that SP9 and DLX factors act as activators in isolation but repress regulatory activity when co-expressed.

Together, these results identify two regulatory modes of SP9: recruitment to enhancers in cooperation with DLX factors, where their combined binding reduces enhancer activity; and direct promoter binding at a smaller subset of sites, including the *Sp9* and *Six3* promoters, where SP9 activates transcription. This is consistent with the directional gene expression changes observed upon *Sp9* perturbation (Figures 2, 3): loss of SP9 would derepress regulatory activity at shared DLX TF-bound enhancers near D1 MSN-associated genes, while removing activation at promoters of key D2 MSN fate drivers such as *Six3*.

### The SP9 ZF domain is required for DLX5 binding and SP9-induced repressive activity

SP family TFs contain a C2H2 ZF domain with three ZFs that typically binds GC-rich motifs (Suske et al., 2005). In osteoblasts, SP7 engages DLX5 via its ZF domain rather than through canonical DNA binding, leading to indirect recruitment to DLX-bound AT-rich motifs (Hojo et al., 2016). To test whether SP9 uses an analogous mechanism, we generated SP9 constructs lacking either the N-terminal domain, which includes the transactivation domain (TAD) (Piskacek et al., 2020) (NΔSP9), or the ZF domain (SP9ΔZF), as well as a construct carrying the patient-associated point mutation p.Glu378Gly within the ZF domain (SP9*378), identified in patients with neurodevelopmental disorders including epilepsy and intellectual disability (Tessarech et al., 2024) (Figure 5A). Co-immunoprecipitation (CoIP) followed by western blotting in HEK293FT cells showed that deletion of the SP9 ZF domain abolished interaction with DLX5 (Figure 5A-C), while mutants lacking the N-domain or carrying the SP9*378 mutation retained DLX5 interaction. Because SP9ΔZF localizes to both the nucleus and cytoplasm (Figure S7A), whole-cell lysates were used for Figure 5C. C2H2-ZFs are increasingly recognized as domains that can regulate transcription not only through DNA binding but also through protein–protein interactions (Brayer and Segal, 2008; DelRosso et al., 2023), consistent with the dual regulatory role of the SP9 ZF domain proposed here.

**Figure 5:**
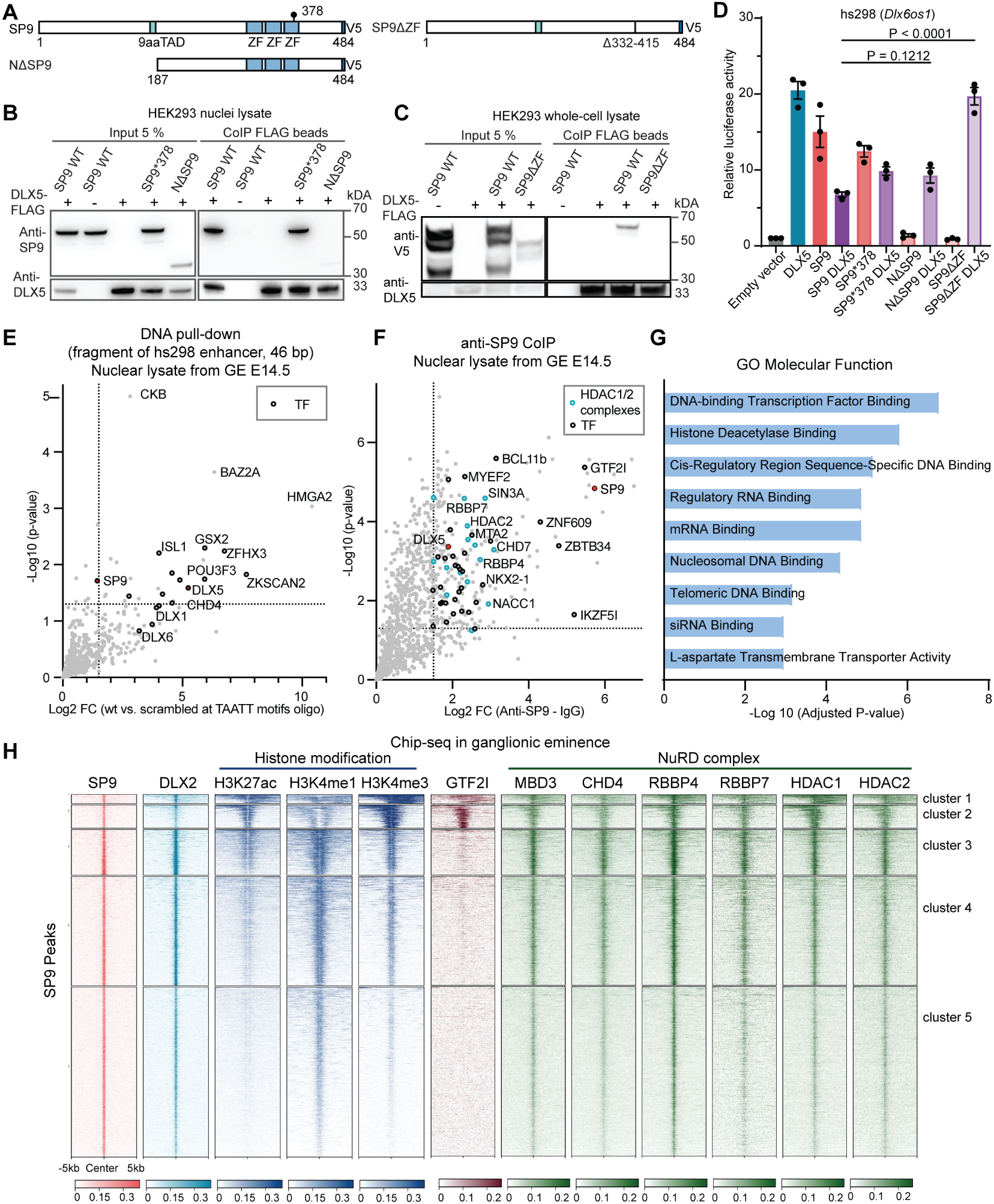
SP9 associates with DLX5 and the NuRD complex in vitro and in the embryonic ganglionic eminence. **A**, Schematic of C-terminally V5-tagged SP9 constructs: wild-type (WT) SP9, an N-terminal deletion mutant (NΔSP9), and a zinc finger (ZF) domain deletion mutant (SP9ΔZF). The nine-amino-acid transactivation domain (9aaTAD), the three ZF domains, and the position of the patient-derived SP9*378 variant are indicated. **B**, Co-immunoprecipitation (CoIP) from nuclear lysates of HEK293FT cells co-transfected with DLX5-FLAG and either SP9-V5 WT, SP9*378-V5, or NΔSP9-V5. Left: input fractions probed with anti-SP9 (top) and anti-DLX5 (bottom). Right: CoIP using anti-FLAG beads, probed with anti-SP9 (top) and anti-DLX5 (bottom). **C**, CoIP from whole-cell lysates of HEK293FT cells co-transfected with DLX5-FLAG and either SP9-V5 WT or SP9ΔZF-V5. Left: input fractions probed with anti-V5 (top) and anti-DLX5 (bottom). Right: CoIP using anti-FLAG beads, probed with anti-V5 (the anti-SP9 epitope lies within the deleted ZF region, requiring V5 detection; top) and anti-DLX5 (bottom). **D**, Luciferase reporter activity driven by the hs298 enhancer of *Dlx6os1* in N2A cells co-transfected with *Dlx5* and either *Sp9*, *Sp9*378*, *N*Δ*Sp9*, or *Sp9*Δ*ZF*. Bars represent mean ± s.e.m. of 9 replicates from 3 independent batches performed in triplicate; points indicate batch means. Statistical significance was assessed by two-way ANOVA with Tukey’s honestly significant difference (HSD) post hoc test. Exact *P*-values are provided in Table S7. **E**, Label-free quantitative mass spectrometry (MS) of proteins captured by DNA pull-down using a 46-bp fragment of the hs298 enhancer of *Dlx6os1* versus the same fragment with the TAATT motifs scrambled, from E14.5 ganglionic eminence (GE) nuclear lysates. Proteins with p ≤ 0.05 and log_2_ LFQ FC ≥ 1.5 were considered significantly enriched (n = 25). Transcription factors (TFs) are indicated in black. **F**, Volcano plot of proteins enriched in anti-SP9 CoIP-MS from E14.5 GE nuclear lysates, relative to IgG controls. Proteins with log_2_FC ≥ 1.5 and *P* ≤ 0.05 were considered significantly enriched (n = 223). Components of HDAC1/2-containing complexes are highlighted in blue; TFs are indicated in black. *P*-values were calculated using a two-sided Student’s t-test with permutation-based false discovery rate (FDR) correction. **G**, Gene Ontology (GO) molecular function enrichment analysis of proteins identified by anti-SP9 CoIP-MS. **H**, Heatmap of ChIP-seq signal intensity across SP9-bound regions for SP9 (Xu et al., 2018; Catta-Preta et al., 2025), DLX2, histone modifications (H3K27ac, H3K4me1, H3K4me3) (Lindtner et al., 2019), GTF2I (Kopp et al., 2020), and NuRD complex subunits (MBD3, CHD4, RBBP4, RBBP7, HDAC1, HDAC2) (Price et al., 2022). Signal is plotted over a ±5 kb window centered on SP9 peak summits and organized by k-means clustering (5 clusters), separating SP9-bound regions by their co-binding patterns with DLX2, NuRD subunits, and active histone marks. All datasets are from mouse E13.5 GE except GTF2I, which is from E13.5 whole brain.

We next assessed the functional consequences of these mutations plus the C-terminal domain mutant (SP9ΔC) in dual-luciferase reporter assays in N2A cells using three enhancers (hs298, hs119, and hs818 near *Pbx3*) and three promoters (*Sp9*, *Six3*, and *Zeb2*) (Figures 5D, S6B-G). Deletion of the SP9 ZF domain abolished SP9-mediated suppression of DLX5-dependent reporter activity, consistent with loss of SP9-DLX5 interaction, although this effect may also reflect reduced nuclear localization. In contrast, NΔSP9 retained the ability to repress DLX5-dependent activity, similar to SP7 (Hojo et al., 2016), but showed reduced activation of REs compared with wild-type (WT) SP9 (Figures 5D, S6C,G), consistent with loss of the TAD in this mutant. The patient mutation SP9*378, which retained DLX5 interaction, showed reduced activation of the GC-rich promoter (Figures 5B, S6D). Together, these results identify two functionally separable domains within SP9 with distinct roles: an N-terminal domain that drives activation at GC-rich promoters, and a ZF domain that mediates both DLX5 interaction and repression of DLX5-dependent activity.

To confirm that SP9 associates with DLX factors at native enhancers *in vivo*, we performed DNA pull-down assays followed by mass spectrometry using a biotinylated 46-bp fragment of the hs298 enhancer (one of the REs used in our reporter assays, containing three TAATT motifs but no canonical SP motifs; Table S8) as bait, with nuclear lysates from E14.5 mouse GE. SP9 and DLX1/5/6 were enriched compared with a control oligonucleotide scrambled at the TAATT motifs (Figure 5E; Table S9), supporting a model in which SP9 occupies TAATT-containing enhancers indirectly via DLX TFs *in vivo*. The pull-down also recovered additional GE-expressed TFs and components of multiple chromatin-modifying complexes with predominantly repressive functions, including NuRD components (Table S9). Several enriched factors are encoded by SFARI ASD-risk genes, including *Chd4*, *Setbp1*, *Ash1l*, *Pou3f3*, and *Ldb1* (Abrahams et al., 2013).

To test whether the functional separation of SP9 modes is preserved across cellular contexts, we performed the luciferase reporter assays in mouse 3T3-L1 cells (Figure S6H-K). In this fibroblast context, SP9 did not activate any of the tested REs but retained the ability to repress DLX5-mediated reporter activity (Figure S6J-K). Thus, activation by SP9 is context-dependent, likely requiring lineage-specific transcriptional cofactors absent in 3T3-L1 cells, whereas repression of DLX-dependent activity was retained in both cellular contexts tested. These findings provide a molecular basis for the two regulatory modes of SP9 identified by genomic and reporter analysis (Figures 3, 4): enhancer-localized repression depends on DLX factor recruitment via the SP9 ZF domain, while promoter-localized activation engages the N-terminal domain.

### SP9 associates with NuRD and DLX factors at repressed enhancers

To identify the endogenous SP9 interactome *in vivo*, we performed co-immunoprecipitation followed by mass spectrometry (CoIP-MS) using SP9-specific antibodies on nuclear lysates from E14.5 mouse GE. We identified 223 proteins enriched over IgG controls (P < 0.05, log2 fold change (FC) > 1.5; Figure 5F; Table S9), including DLX1/5/6, multiple TFs involved in GABAergic neuron development (BCL11B, NKX2-1, PBX3, TCF4/12, SATB2), and GTF2I, a general TF previously implicated in neuronal maturation (López-Tobón et al., 2023). Gene Ontology (GO) molecular function analysis revealed enrichment for histone deacetylase (HDAC) binding (Figure 5G), with hits including HDAC1/2, MTA1/2, RCOR1, RBBP4/7, and SIN3A, which are components of multiple HDAC chromatin-regulatory complexes including NuRD, SIN3, and CoREST (Alshehri et al., 2025; Nitarska et al., 2016).

To resolve which of these complexes SP9 associates with, we performed reciprocal CoIP-MS using anti-HDAC2 and anti-SIN3A antibodies on E14.5 GE nuclear lysates. HDAC2 co-immunoprecipitates enriched SP9 and DLX2/5 along with canonical NuRD-specific components (MTA1/2, MBD3, CHD4, RBBP4/7), whereas SIN3A co-immunoprecipitates did not recover SP9 or DLX2/5 (Figure 5L–M; Table S9), indicating that SP9 associates with HDAC1/2 through the NuRD complex rather than SIN3. This is consistent with previous work showing that DLX1 interacts with NuRD components in the GE (Price et al., 2022).

To distinguish direct protein-protein interaction from interactions bridged by shared nucleic acids, we treated N2A nuclear lysates (transfected with *Sp9*-V5) with the endonuclease Benzonase prior to SP9-V5 CoIP-MS. NuRD components, including HDAC1 and HDAC2, were still recovered (Table S9), indicating that the interaction is independent of DNA or RNA.

To map the SP9 domains required for these interactions, we performed CoIP-MS in N2A cells transfected with *Sp9*-V5, *N*Δ*Sp9*-V5, or *Sp9*Δ*C*-V5 (Figure S6N, Table S9). NΔSP9 showed reduced interaction with coactivators including KAT2A and EP300 and failed to activate reporters in luciferase assays (Figures 5D, S6C-G, N), whereas SP9ΔC displayed an interactome comparable to full-length SP9. Consistent with the functional separation of SP9 domains shown in Figure 5, the N-terminal domain is required for coactivator recruitment and transcriptional activation, whereas interactions with DLX factors and NuRD components depend on the zinc-finger domain and not the C terminus.

To examine the genomic distribution of interactors identified in SP9 CoIP-MS from the GE, we analyzed published ChIP-seq datasets for DLX2 along with histone modifications (H3K27ac, H3K4me1, H3K4me3) (Lindtner et al., 2019), GTF2I (Kopp et al., 2020), and NuRD components (MBD3, CHD4, RBBP4, RBBP7, HDAC1, HDAC2) (Price et al., 2022).

Unsupervised clustering of SP9 peaks based on these signals identified promoter-like regions (clusters 1 and 2) marked by H3K4me3 and H3K27ac and distal regions (clusters 3-5) with lower H3K27ac and H3K4me1 (Figure 5H). GTF2I co-occupied a subset of promoter-like SP9 peaks, consistent with its preference for GC-rich KLF/SP motifs (Kopp et al., 2020). NuRD components were detected across both promoter-like and distal SP9 peaks. A subset of distal SP9 peaks with strong SP9 and DLX2 co-binding retained H3K4me1 but showed reduced H3K27ac (Figure 5H), consistent with repressed enhancers and with the SP9+DLX-mediated repression in Figure 4L-O.

Together, these results identify SP9 as a partner of DLX TFs and NuRD chromatin regulators *in vivo*, with the SP9 N-terminal domain mediating coactivator recruitment and the ZF domain mediating DLX interaction, providing a molecular basis for the SP9+DLX-dependent enhancer repression observed in Figure 4L-O.

### The SP9-to-DLX5 ratio redirects SP9 binding between regulatory modes

The genomic and proteomic data above show that SP9 and DLX factors share REs and exist within common protein complexes, indicating they may act combinatorially in gene regulation. To test whether SP9 occupancy depends on DLX5 levels, we overexpressed C-terminally V5-tagged SP9 and C-terminally FLAG-tagged DLX5 in N2A cells, alone or in combination, and profiled binding by cleavage under targets and release using nuclease (CUT&RUN) using anti-V5 and anti-FLAG antibodies (OE1; Figure 6A-B). SP9 peaks identified from the SP9-only and SP9+DLX5 conditions were merged and partitioned into four clusters by k-means clustering based on their relative SP9 signal across both conditions (Figure 6C-D). Three clusters (C1-C3) showed reduced SP9 occupancy upon DLX5 coexpression, while a fourth cluster (C4) showed increased SP9 occupancy.

**Figure 6:**
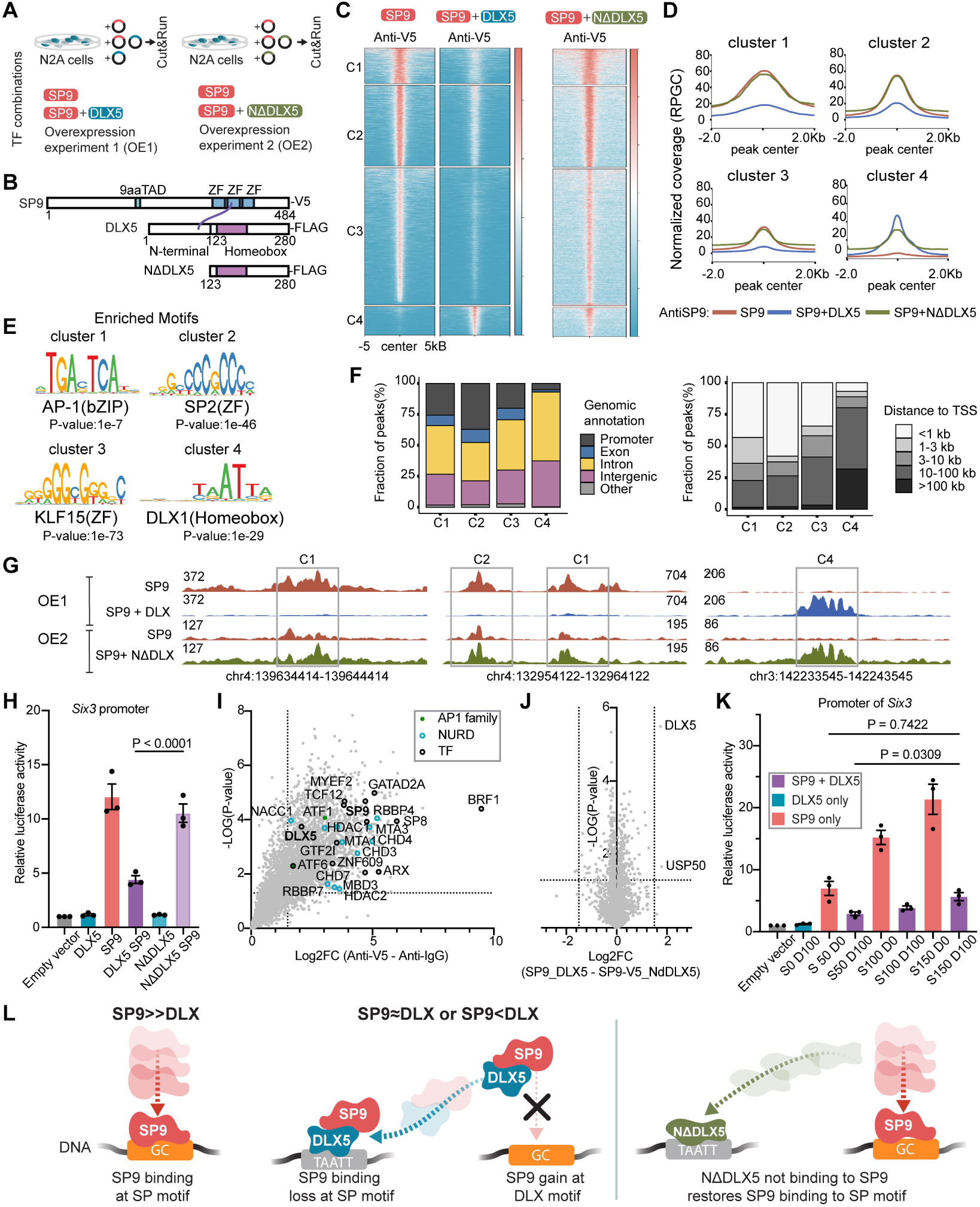
SP9 genomic occupancy is redistributed from SP motifs toward DLX motifs in the presence of DLX5. **A**, Schematic of the transcription factor (TF) overexpression strategy in N2A cells. Cells were transfected with SP9 alone or co-expressed with WT DLX5 (overexpression experiment 1, OE1) or with the N-terminal deletion mutant NΔDLX5 (OE2), followed by CUT&RUN profiling. **B**, Domain organization of SP9, DLX5, and NΔDLX5, highlighting the zinc finger (ZF) and homeobox domains. **C**, Heatmaps of SP9 CUT&RUN signal (anti-V5) centered on SP9 peak regions (±5 kb), clustered into four groups (C1-C4) by k-means. Conditions: SP9 alone, SP9+DLX5 (OE1), SP9+NΔDLX5 (OE2). **D**, Aggregate signal plots for each cluster (C1-C4) showing normalized SP9 binding profiles at peak centers under the three conditions in (C). **E**, Top enriched motifs identified by motif analysis of peaks in each cluster, with associated *P*-values. **F**, Genomic annotation of SP9-bound peaks across clusters. Left: distribution across promoter, exon, intron, intergenic, and other regions. Right: distance to transcription start sites (TSS). **G**, Genome browser tracks at three representative loci showing SP9 binding under OE1 and OE2 conditions, illustrating loss of SP9 occupancy upon DLX5 coexpression at SP-motif-containing sites and rescue with NΔDLX5, as well as gain of SP9 occupancy at DLX-motif-containing sites with WT DLX5. **H**, Luciferase reporter activity driven by the *Six3* promoter in N2A cells transfected with combinations of *Sp9*, *Dlx5*, and *N*Δ*Dlx5* as indicated. **I**, Volcano plot of co-immunoprecipitation followed by mass spectrometry (CoIP-MS) using anti-V5 in N2A cells co-transfected with *Sp9-V5* and *Dlx5-FLAG*, relative to IgG controls. Proteins with log_2_FC > 1.5 and *P* < 0.05 were considered significantly enriched (n = 2254). NuRD complex components are highlighted in blue, transcription factors (TFs) in black, and AP-1 family TFs in green. **J**, Volcano plot comparing the anti-V5 CoIP-MS interactomes between SP9+DLX5 and SP9+NΔDLX5 conditions in N2A cells, using the same cutoffs as in (I). **I,J**, *P*-values were calculated using a two-sided Student’s t-test with permutation-based false discovery rate (FDR) correction. **K**, Luciferase reporter activity driven by the *Six3* promoter in N2A cells transfected with varying amounts (ng plasmid DNA) of SP9 (S) and DLX5 (D) to assess ratio-dependent transcriptional activation. **H-K**, Bars represent mean ± s.e.m. of 9 replicates from 3 independent batches performed in triplicate; points indicate batch means. Statistical significance was assessed by two-way ANOVA with Tukey’s honestly significant difference (HSD) post hoc test. Exact *P*-values are provided in Table S7. **L**, Proposed mechanism. When SP9 levels exceed DLX (SP9≫DLX), SP9 binds directly at GC-rich SP motifs. When SP9 and DLX are comparable or DLX is in excess (SP9≈DLX or SP9<DLX), DLX5 sequesters SP9 to TAATT-containing DLX motifs through the DLX5 N-terminal domain, away from its direct SP-motif binding sites. NΔDLX5, which cannot interact with SP9, fails to redirect SP9 to DLX motifs, restoring SP9 binding at SP motifs.

To characterize where these opposing effects occurred, we performed motif enrichment (Figure 6E) and genomic annotation (Figure 6F) on the two sets of regions. The clusters with reduced SP9 binding (C1-C3) were enriched for the AP-1 motif (bZIP family) and the GC-rich SP/KLF motifs (SP2, KLF15) and were predominantly located at promoter and proximal regulatory regions. In contrast, C4, where SP9 occupancy increased, was enriched for the DLX homeobox motif and was predominantly distal. Thus, DLX5 coexpression redirects SP9 binding from SP/KLF- and AP-1-motif-containing proximal REs to DLX-motif-containing distal REs (Figure 6G).

To test whether direct SP9-DLX5 interaction underlies this redirection, we generated an N-terminal deletion mutant of DLX5 (NΔDLX5), based on previous work showing that DLX5 engages SP7 via its N-terminal domain (Hojo et al., 2016). In N2A cell CoIP-MS experiments, NΔDLX5 failed to interact with SP9, without altering the composition of SP9-associated protein complexes (Figure 6I-J), confirming that the N-terminal domain of DLX5 mediates the SP9-DLX5 interaction. We then asked whether loss of this interaction abolishes the DLX5-dependent redirection of SP9 occupancy. Repeating CUT&RUN profiling under coexpression with NΔDLX5 (OE2; Figure 6A-B), NΔDLX5 failed to reduce SP9 occupancy at regions where WT DLX5 caused loss of SP9 binding, with SP9 levels remaining comparable to the SP9-only condition (Figure 6C-D). Profiling of NΔDLX5 binding by CUT&RUN showed that the mutant occupied the same genomic regions as wild-type DLX5 (Figure S7C), excluding a general loss of DLX5 binding. Consistent with these results, NΔDLX5 did not inhibit SP9-driven activation of the *Six3* and *Sp9* promoters in dual-luciferase reporter assays (Figures 6H, S7D). The N-terminal domain of DLX5 is therefore required for both SP9-DLX5 interaction and the DLX5-dependent redirection of SP9 occupancy. Residual SP9 binding at distal DLX-motif sites (cluster C4) under NΔDLX5 coexpression suggests that recruitment to these sites is not exclusively mediated by direct SP9-DLX5 contact; indirect tethering or chromatin-accessibility effects independent of the DLX5 N-terminal domain, or a pioneer-like function of DLX5, may contribute.

To examine whether the SP9-DLX5 balance determines SP9 transcriptional output in a ratio-dependent manner, we performed dual-luciferase reporter assays in N2A cells using the *Six3* promoter and the hs170 enhancer of *Fign*, varying the ratio of SP9 to DLX5 plasmid amounts (Figures 6K, S7E). Reporter activity at the GC-rich *Six3* promoter required SP9 expression to exceed DLX5 expression; at low SP9/DLX5 ratios, SP9 activation was reduced (Figure 6L).

Finally, to assess the *in vivo* relevance of this ratio-dependent mechanism, we examined *Sp9* and *Dlx1/2/5/6* expression patterns in the gLacZ control subset of our single-cell transcriptomic data. In progenitor populations, *Sp9* expression overlapped with *Dlx1/2/5/6* expression. Along the projection neuron pseudotime trajectory, the ratio of *Sp9* to summed *Dlx1/2/5/6* expression increased in a subset of postmitotic cells preceding the D1/D2 MSN bifurcation, and was further elevated specifically in the D2 MSN population (Figure 7A-B).

**Figure 7:**
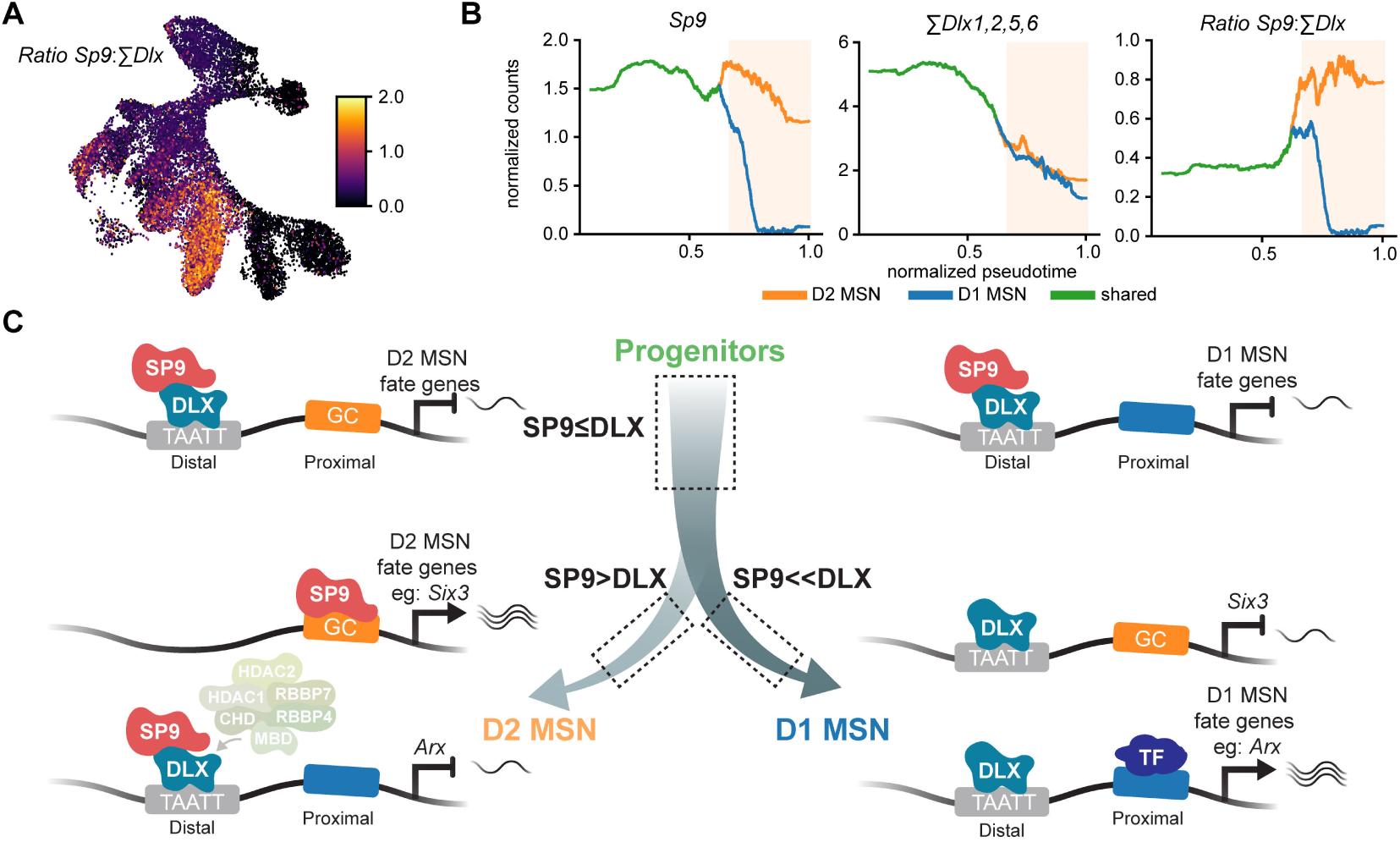
Stoichiometric SP9-DLX factor partnerships specify D1 versus D2 MSNs fate. **A**, UMAP visualization of the GABAergic single-cell transcriptomic dataset, colored by the per-cell ratio of normalized *Sp9* expression counts to the summed expression of *Dlx1, Dlx2, Dlx5*, and *Dlx6* (Ratio *Sp9*:Σ*Dlx*). A subset of cells in the shared trunk before the D1-D2 MSN bifurcation already shows elevated ratios, possibly representing cells biased toward the D2 MSN fate. **B**, Expression dynamics along the projection neuron pseudotime trajectory for *Sp9* (left), summed *Dlx1/2/5/6* expression (middle), and their ratio *Sp9*:Σ*Dlx* (right). Lines indicate the D1 MSN trajectory (blue), the D2 MSN trajectory (orange), and the shared trajectory of progenitors and early postmitotic precursors (green). Shaded region marks the postmitotic branch following the D1-D2 MSN bifurcation. **C**, Proposed model of SP9-DLX stoichiometry in MSN fate control. In progenitors and early postmitotic precursors (SP9≤DLX), SP9-DLX factor complexes occupy DLX-motif-containing distal regulatory elements (REs) of D1 and D2 MSN fate genes; GC-rich promoters of D2 MSN genes such as *Six3* remain largely unbound by SP9. Along the D2 MSN trajectory (SP9>DLX), the relative increase in SP9 enables two simultaneous effects: SP9 binding at the GC-rich *Six3* promoter promotes D2 MSN-fate gene expression, while SP9-DLX factor complexes at distal D1 MSN-gene REs recruit the NuRD complex (CHD, MBD, RBBP4/7, HDAC1/2) to repress D1 MSN-fate genes. Along the D1 MSN trajectory (SP9<<DLX), SP9 is absent from these REs; DLX factors remain bound at distal TAATT sites, and D1 MSN-fate genes such as *Arx* are activated through other transcription factors (TFs) at their proximal REs.

Together, these results support a mechanism in which the relative abundance of SP9 and DLX5 at shared REs determines SP9 binding distribution between SP-motif promoters and DLX-motif distal REs (Figure 7C), providing a molecular basis for the lineage redirection observed upon *Sp9* loss (Figures 1-3).

## Discussion

Here, we show that SP9 and DLX factors interact at shared REs to control lineage allocation during GABAergic neuron development in the GE. Sparse *in vivo* CRISPR perturbation combined with clonal lineage tracing and single-cell transcriptomics showed that loss of *Sp9* does not eliminate the LGE neuronal lineage but redirects individual progenitors from D2 MSN fate toward D1 MSN and ITC fates. Mechanistically, SP9 operates through two regulatory modes that depend on distinct functions of its ZF domain. At TAATT-containing distal REs, SP9 is recruited through ZF-dependent protein-protein interaction with DLX factors, and their combined occupancy represses regulatory activity. At a smaller set of GC-rich promoters, including *Six3* and *Sp9* itself, SP9 directly binds DNA through its ZF domain and activates transcription through its N-terminal domain. These findings support a stoichiometric switch model in which the relative abundance of SP9 and DLX factors, reinforced by SP9 self-activation and patterned by differential TF expression across individual cells, balances D1 and D2 MSN fate allocation (Figure 7C).

Our findings build on a sequence of prior loss- and gain-of-function studies that collectively position SP9 as a key determinant of D2 versus D1 MSN identity (Zhang et al., 2016; Xu et al., 2018; Li et al., 2022). *Sp9*-null mice lose the majority of D2 MSNs while D1 MSN development remains largely unaffected, and conversely, *Sp9* overexpression in LGE progenitors gradually converts pre-D1 cells into D2-like MSNs that survive normally into adulthood. Our data confirm this framework, additionally identify ITC cells as an alternative *Sp9*-dependent fate, and provide a mechanistic basis for the switch (Figure 3J).

Our approach, combining CRISPR perturbation with lineage barcoding and clonal-state embedding, addresses two limitations in the study of neuronal fate specification. First, conditional knockout strategies perturb all progenitors simultaneously and lose information about the cellular trajectories of individual perturbed cells, making it difficult to separate effects intrinsic to the perturbed cells from effects mediated by the broader tissue context. Second, prior single-cell lineage analyses have relied on coupling between discrete transcriptomic clusters (Bandler et al., 2022; Wagner et al., 2018), which forces lineage analysis into predefined pairwise categories and obscures multipotency and graded fate biases. By coupling CRISPR perturbation in a sparse subset of GE progenitors with TrackerSeq (Bandler et al., 2022; Dvoretskova et al., 2024) and clone2vec embedding (Erickson et al., 2025; Isaev et al., 2026), we resolved continuous spectra of clonal fate biases and the cellular consequences of perturbation in the same dataset. This combination showed that *Sp9* loss redirects clonal output along a continuum of fate biases (Figure 3). We anticipate that this strategy can be applied broadly to dissect cell-autonomous contributions of fate-determining TFs during *in vivo* development.

At the molecular level, SP9 operates through two functionally distinct regulatory modes, distinguished by genomic context, partner availability, and direction of transcriptional regulation (activation vs. repression). At the majority of SP9-bound regions (distal, DLX-motif-containing enhancers), SP9 binding depends on its ZF-mediated interaction with DLX factors rather than on direct DNA contact. CoIP-MS from GE nuclear lysates identified NuRD chromatin regulators (HDAC1/2, MTA1/2, MBD3, CHD4, RBBP4/7) among SP9-DLX-associated proteins (Figure 5F). Recruitment of NuRD and its associated deacetylase activity could contribute to the reduced regulatory activity observed at SP9–DLX co-bound regions in reporter assays (Koulle et al., 2026; Alshehri et al., 2025; Asmamaw et al., 2024). In contrast, at a smaller set of promoters with GC-rich SP motifs, SP9 directly binds DNA through its ZF domain and activates transcription through its N-terminal domain, a function lost when the N-terminal domain is removed. A similar domain architecture has been described for SP7 in osteoblasts, where its N-terminal domain counteracts the repressive output of its ZF-mediated interaction with DLX5 (Hojo et al., 2016), suggesting that the activation-repression balance is set by the relative contributions of these two domains within a single SP-family protein. The SP9 ZF domain thus fits a broader paradigm in which C2H2 ZF domains engage both DNA and partner proteins (Brayer and Segal, 2008; DelRosso et al., 2023). The principle that direct TF interactions reshape each other’s genomic occupancy, as seen here for SP9 and DLX, has been described in other developmental systems, including TBX5, NKX2-5, and GATA4 in the heart (Luna-Zurita et al., 2016) and NGN2 and EMX1 in the developing nervous system (Ang et al., 2024). These examples support a broader paradigm in which protein-protein interactions specify lineage-specific regulatory programs beyond what DNA-binding motifs alone predict.

Beyond this mechanistic dichotomy, the relative abundance of SP9 and DLX factors, rather than their individual levels, determines SP9 occupancy distribution between the two modes. Reporter assays at the GC-rich *Six3* promoter showed ratio-dependent activation that required SP9 levels to exceed DLX5; at low SP9/DLX5 ratios, SP9 was unable to drive promoter activation (Figure 6K). Among SP9’s direct promoter targets is *Six3*, a known master regulator of D2 MSN fate (Song et al., 2021), suggesting that the broad downregulation of D2 MSN markers upon *Sp9* loss is likely amplified through loss of SIX3-dependent transcription.

The redistribution of SP9 binding toward distal DLX-motif sites upon DLX5 coexpression indicates a competitive mechanism in which DLX factors sequester SP9 to TAATT-containing distal REs, limiting its occupancy at GC-rich promoter sites. This sequestration depends on the DLX5 N-terminal domain containing its intrinsically disordered region, which engages SP9 directly: deletion of the N-terminus disrupts SP9-DLX5 interaction and abolishes the redistribution of SP9 binding observed with full-length DLX5 (Figure 6L). The competition has a built-in amplification: among SP9’s direct promoter targets is the *Sp9* gene itself, which in our reporter assays was activated by SP9 protein. Once SP9 protein rises sufficiently in individual cells to outcompete DLX-mediated sequestration, self-activation reinforces *Sp9* transcription and stabilizes the D2 MSN trajectory (Figure 7C). *In vivo*, single-cell expression patterns along pseudotime trajectories are consistent with this model. *Sp9* and *Dlx1/2/5/6* expression overlap broadly in progenitors. As cells progress along the postmitotic D2 MSN trajectory, *Sp9* expression rises sharply in a subset of cells, while *Dlx1/2/5/6* expression decreases similarly in both the D1 and D2 MSN trajectories (Figure 7A,B). This selective rise in *Sp9*, combined with the parallel decline of *Dlx* TFs in both trajectories, increases the *Sp9*-to-*Dlx* TFs ratio specifically in the D2 MSN branch. Combined, the reporter titrations and *in vivo* expression patterns support a stoichiometric switch model in which the developmental tuning of cooperating TF levels translates continuous variation in TF concentration into discrete fate. The complementary question of how *Sp9* expression is actively downregulated along the D1 MSN trajectory remains open for future work.

Recent integrative analyses of human and mouse brain development implicate GE-derived neurons in ASD, intellectual disability, and other neurodevelopmental conditions (Aivazidis et al., 2025; Satterstrom et al., 2020; Evans et al., 2024), and disease-risk genes are disproportionately expressed in GE progenitors and their immediate progeny. SP9 itself sits at this intersection: Clinical variant databases and patient studies indicate that *SP9* variants, including stop-gain and missense alleles, are associated with NDDs, with clinical features including epilepsy and intellectual disability (Tessarech et al., 2024; Landrum et al., 2018). Tessarech et al. identified three patients with *SP9* mutations in the ZF domain, including the p.Glu378Gly variant examined here (SP9*378). In our assays, SP9*378 retained interaction with DLX factors but showed reduced activation of GC-rich promoters in N2A cells (Figure 5B; Figure S6D), suggesting that disease-associated mutations in the ZF domain can selectively impair the SP9 promoter-activation mode while leaving SP9-DLX association intact. More broadly, *Sp9* loss deregulated several epilepsy-associated genes in IN populations (*Gria1*, *Kcnd2*, *Nrxn1*, and *Cacna1c*; (Piacentini et al., 2026); Figure S3), and SP9 interactors in the GE included multiple SFARI ASD-associated proteins, such as BCL11B, CHD7, SATB2, TCF20 and GTF2I (Abrahams et al., 2013) (Table S9). These observations link SP9 function to a broader set of GE-derived NDD genes and raise the possibility that distinct *SP9* mutations differentially affect the regulatory modes identified here.

The stoichiometric partnership described here for SP9 and DLX factors is likely embedded within a broader context-dependent regulatory network in the GE. Our CoIP-MS analyses identified additional GABAergic lineage-specific TFs among SP9 interactors, including NKX2-1, PBX3, TCF12, and BCL11B (Figure 5F; Table S9). Among the four DLX family members expressed in the GE (DLX1, DLX2, DLX5, DLX6) (Eisenstat et al., 1999; Lindtner et al., 2019; Rubenstein et al., 2024), DLX2 acts as a stronger activator while DLX1 functions more prominently as a repressor (Lindtner et al., 2019; Dvoretskova et al., 2024; Price et al., 2022), consistent with the differential reporter activities we observed at SP9+DLX co-bound REs (Figure 4L-O). Context-dependent partnerships have been described for SP7, which engages distinct TFs across osteoblasts, osteocytes, and oligodendrocytes (Göös et al., 2022; Hojo et al., 2016; Wang et al., 2021; Elbaz et al., 2024), suggesting that the specific complement of available cofactors may similarly shape SP9 activity across the different cell states in which it is expressed. Differences in SP9-mediated activation between N2A and 3T3-L1 cells in our assays may therefore reflect differential cofactor availability (Wu et al., 2016; Frömel et al., 2025). Beyond DLX factors, MEIS2 likely operates within this framework: it co-binds at shared cis-regulatory elements (CREs) with DLX5 (Dvoretskova et al., 2024) and loses predictive value for fate upon *Sp9* perturbation in our CatBoost analysis (Figure 3K), although whether it cooperates with or competes against SP9 for DLX partners remains an open question. A second candidate partnership involves AP-1 family TFs: AP-1 motifs were enriched at a subset of SP9 binding sites lost upon DLX5 coexpression, AP-1 family TFs were recovered in N2A SP9-V5 CoIP-MS as well (Figure 6C-E,I; Table S9), and SP7 has been reported to engage AP-1 motifs in osteocytes (Wang et al., 2021). Whether SP9 similarly partners with AP-1 factors in the GE remains to be tested; such indirect cofactor recruitment could in principle account for activation of enhancers lacking SP motifs by SP9 alone in N2A cells (Figure 4M-N).

### Limitations of the study

Several limitations of this study should be acknowledged. First, although our data support a model in which loss of SP9 activation of *Six3* accounts for much of the broad D2 MSN marker downregulation, we cannot rule out that additional D2 MSN markers are direct SP9 targets; available SP9 ChIP-seq data (Xu et al., 2018; Catta-Preta et al., 2025) derive from bulk E13.5 GE rather than cell-state-specific populations, precluding resolution of binding patterns within emerging D2 versus D1 MSN lineages. Second, we did not address whether the patient mutation SP9*378 disrupts SP9 association with NuRD. Third, CRISPR-induced frame-shift indels produce heterogeneous editing outcomes across perturbed cells, and we did not perform per-cell genotyping or genetic rescue experiments to formally validate biallelic *Sp9* loss or exclude off-target contributions. Fourth, our IUE strategy under-targeted MGE-derived populations relative to LGE-derived populations, in particular *Lhx6*^+^ INs and *Lhx8*^+^ cholinergic precursors, which were inconsistently recovered across replicates; our dataset is therefore underpowered to test whether *Sp9* loss alters *Lhx6*^+^ interneuron or cholinergic neuron specification.

Beyond these caveats, stoichiometric TF partnerships may represent a quantitative principle of fate allocation that operates wherever related cell types arise from progenitors sharing a common TF repertoire, with direct implications for NDDs that converge on GE development. How this initial asymmetry in TF stoichiometry is established among sister progenitors remains the central open question.

## Supplementary Figures

**Figure S 1:**
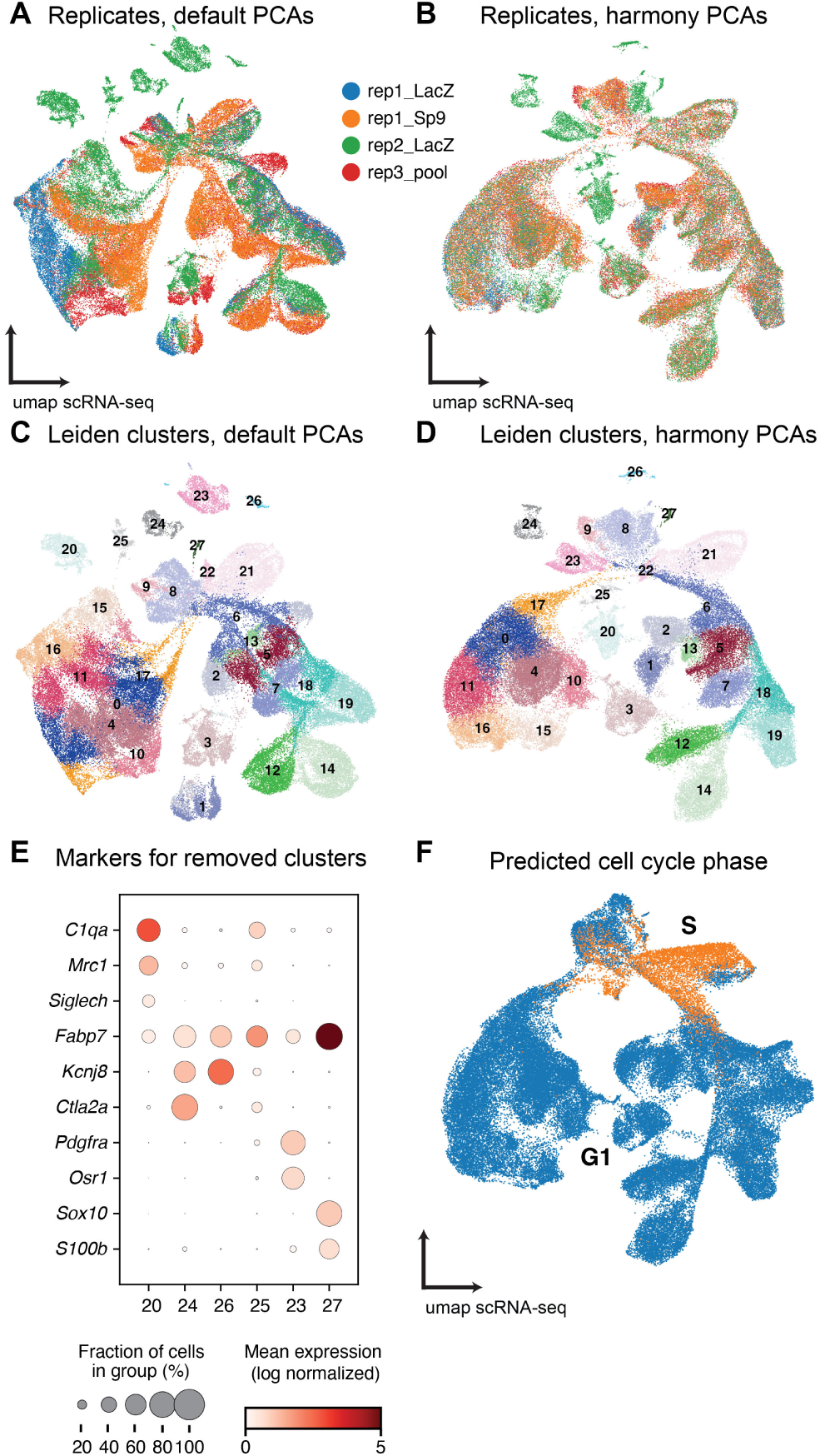
Preprocessing of the scRNA-seq tCROP-seq dataset from E16.5 mouse forebrain, related to Figure 1. **A**, UMAP of the full perturbation dataset using standard Scanpy parameters, colored by sample. The sample labeled “rep3_pool” combines cell suspensions from multiple embryos pooled before scRNA-seq library preparation; each contributing embryo received a single gRNA type (gLacZ or gSp9). This sample serves as an internal control for batch versus perturbation effects. **B**, UMAP of the same dataset after Harmony batch correction of principal components. **C**, UMAP using standard Scanpy parameters, colored by initial Leiden clusters at resolution 1.0. **D**, UMAP after Harmony batch correction, showing the same Leiden clusters as in (C). **E**, Dot plot of vascular and immune cell-type marker expression in initial Leiden clusters that were subsequently removed from the dataset. **F**, UMAP colored by predicted cell-cycle phase, inferred using sc.tl.score_genes_cell_cycle() with mouse-ortholog gene lists.

**Figure S 2:**
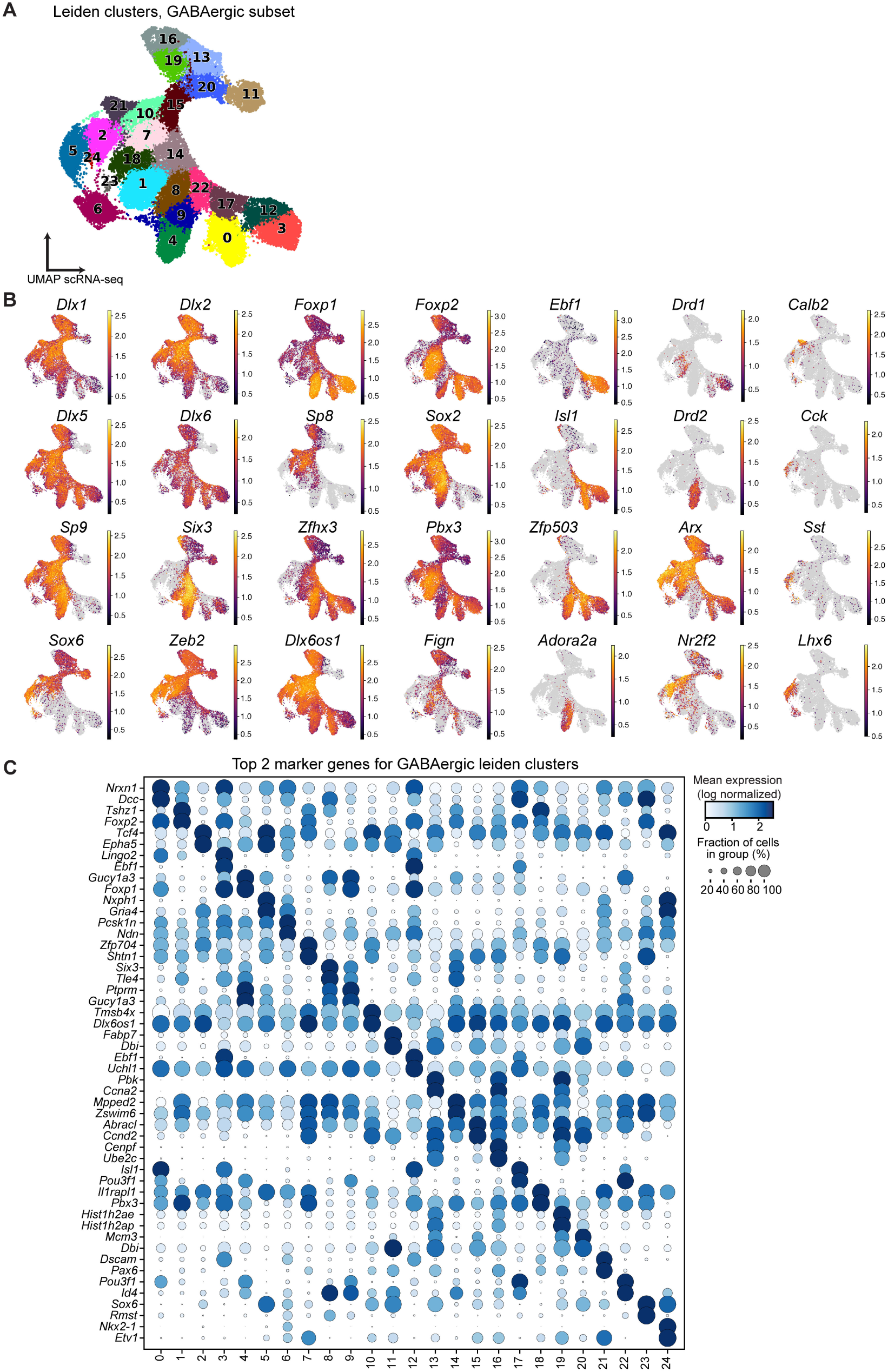
GABAergic marker gene expression supporting the GABAergic cell subset of the scRNA-seq dataset, related to Figure 2. **A**, UMAP of the GABAergic scRNA-seq subset, colored by Leiden clusters at resolution 1.5. The same subset is shown in Figure 2A with cell-type annotations, and the same clusters appear on the PAGA trajectories in Figure 2C. **B**, Feature plots of selected GABAergic marker genes in the GABAergic scRNA-seq subset, restricted to gLacZ control cells. Counts are scaled and log-normalized. **C**, Dot plot of the top 2 marker genes per Leiden cluster, computed from gLacZ control cells only. Mean expression is log-normalized between 0 and 1 across all cells in each cluster.

**Figure S 3:**
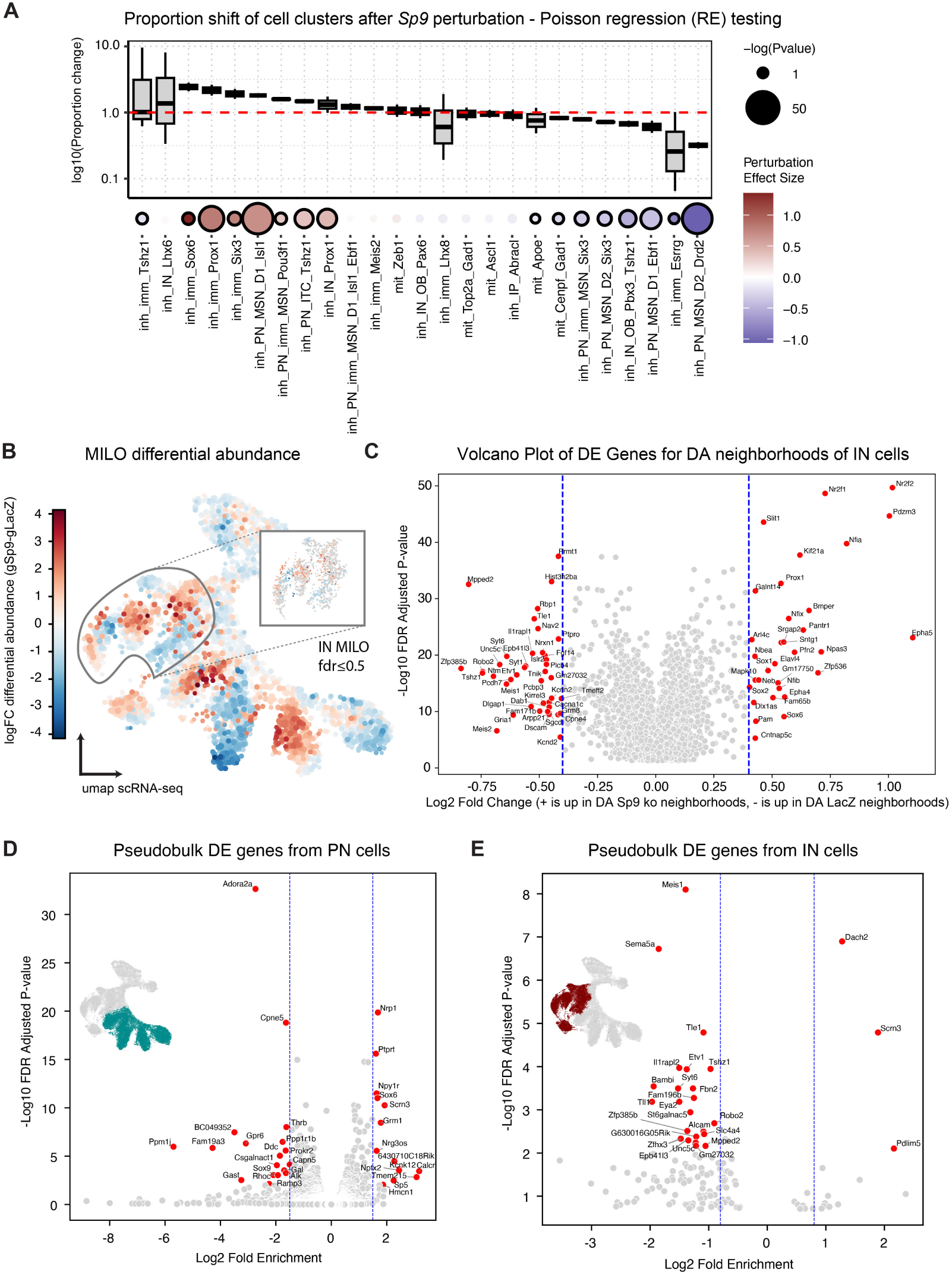
Differential abundance analysis of *Sp9* perturbation, related to Figure 2. **A**, Top: relative change in cell numbers in gSp9 compared to gLacZ control across inhibitory neuron clusters. Y-axis: log_10_ ratio of (proportion of cells of the given type in gSp9-perturbed embryos) over (proportion of cells of the given type in gLacZ controls). Bottom: perturbation effect per cluster. Dot color corresponds to effect size, dot size corresponds to − log_10_(*P*-value). *P*-values were derived from Poisson regression (RE) models with FDR correction. Black outline around a dot indicates statistical significance (*P* ≤ 0.01). Box plots show the median (center line), quartiles (box bounds), and 1.5 × interquartile range (whiskers). **B**, Differential abundance (DA) testing (MILO; knn = 100) across mitotic and GABAergic cells. Neighborhoods enriched for gSp9 cells compared to gLacZ cells are shown in red, depleted neighborhoods in blue. The outlined region contains the interneuron (IN) neighborhoods used for the differential expression (DE) analysis in (C). Inset shows neighborhoods meeting a false discovery rate (FDR) threshold of ≤ 0.5, a higher FDR threshold was selected compared to the projection neuron (PN) subset since abundance shifts in the IN subset had lower absolute magnitudes. **C**, Volcano plot of DE genes between significantly enriched and depleted IN neighborhoods. Genes meeting | log_2_FC| ≥ 2 and FDR ≤ 0.5 are highlighted in red. **D**, Per-sample pseudobulk differential gene expression analysis (PyDESeq2) of PNs and their precursor states between gSp9 and gLacZ conditions. Highlighted points: | log_2_ FC| ≥ 1.5 and adjusted *P* ≤ 0.01. Inset shows on the GABAergic UMAP the cells included in this analysis. **E**, PyDESeq2 of INs and their precursor states between gSp9 and gLacZ conditions. Highlighted points: | log_2_ FC| ≥ 0.8 and adjusted *P* ≤ 0.01. Inset shows on the GABAergic UMAP the cells included in this analysis.

**Figure S 4:**
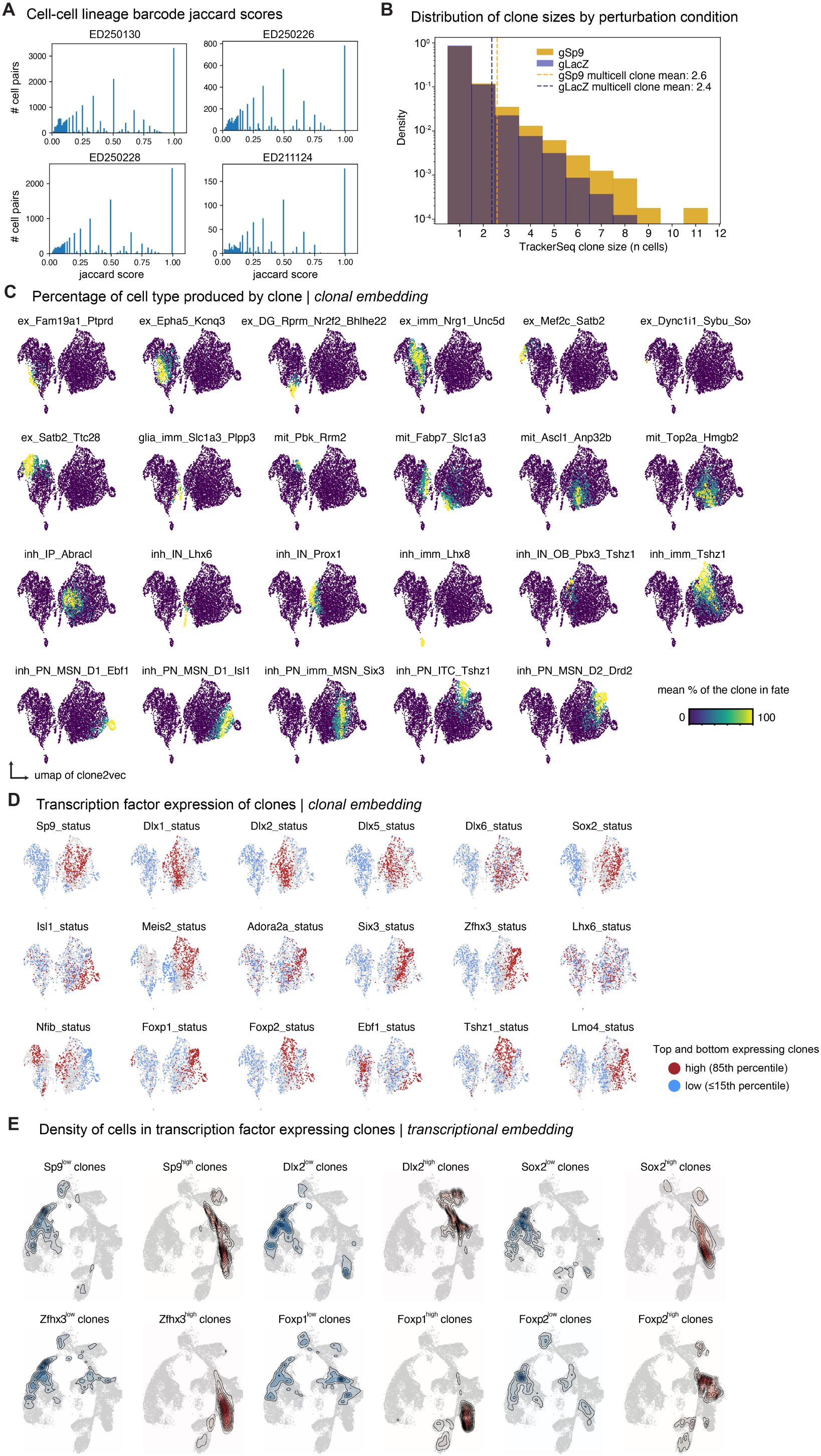
Clone-size distributions, per-clone cell-type output, and gene expression of TrackerSeq clones, related to Figure 3. **A**, Distribution of Jaccard scores of TrackerSeq lineage barcodes for each cell-cell pair according to the shared pattern of lineage barcodes detected in each cell. Edges ≤0.5 were removed for all libraries included in this study to ensure high quality clonal information for analysis. **B**, Histogram of TrackerSeq clone sizes split by perturbation condition (gLacZ, gSp9). Dashed vertical lines indicate the mean clone size of multi-cell clones in each condition (gSp9: 2.6; gLacZ: 2.4). **C**, Clone2vec UMAPs colored by the mean percentage of cells per clone classified into each annotated cell type (extending Figure 3F, which shows the four-class summary). Color scale: 0-100% mean clone composition. **D**, Clone2vec UMAPs colored by per-clone aggregated expression of selected GABAergic differentiation transcription factors (TFs). Clones in the top 15% (high, red, ≥85th percentile) and bottom 15% (low, blue, ≤15th percentile) of aggregate expression are highlighted; remaining clones are shown in grey. **E**, Kernel density plots on the scRNA-seq UMAP showing the distribution of member cells from clones in the top 15% (high; red density) or bottom 15% (low; blue density) of per-clone aggregated TF expression, for selected PN-fate-associated TFs. The differing density distributions between high- and low-expressing clones visualize how TF expression level in a clone predicts the cell-type composition of its member cells.

**Figure S 5:**
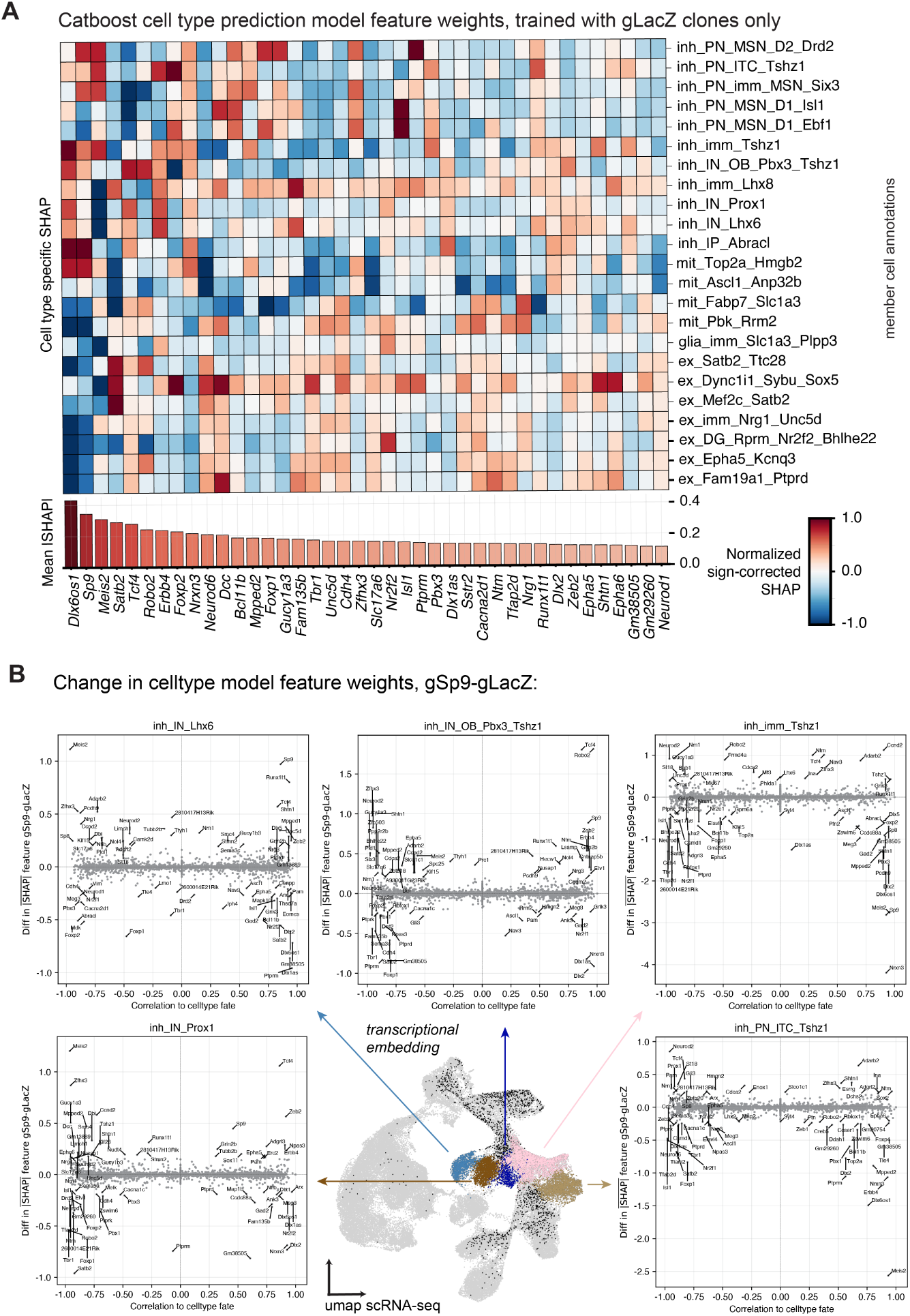
Lineage data-trained CatBoost cell type prediction model, related to Figure 3. **A**, Top gene SHAP values from CatBoost cell-type prediction model. Global SHAPs are shown in rank order at the bottom with cell-type specific SHAPs shown in the heatmap. Positive feature weights indicate that gene expression predicts cell type output, while negative values are antipredictive of the cell fate shown. **B**, Comparison of CatBoost cell-type prediction model gene SHAP values produced for predictions of specific cell states. Correlation of gene features with the cell fate are on the x-axis, the change in feature weight after SP9 perturbation is shown on the y-axis. gLacZ model SHAPs were subtracted from gSp9 SHAPs. All cells belonging to clones which produced cells in any of the highlighted clusters are shown in black on the scRNA-seq UMAP.

**Figure S 6:**
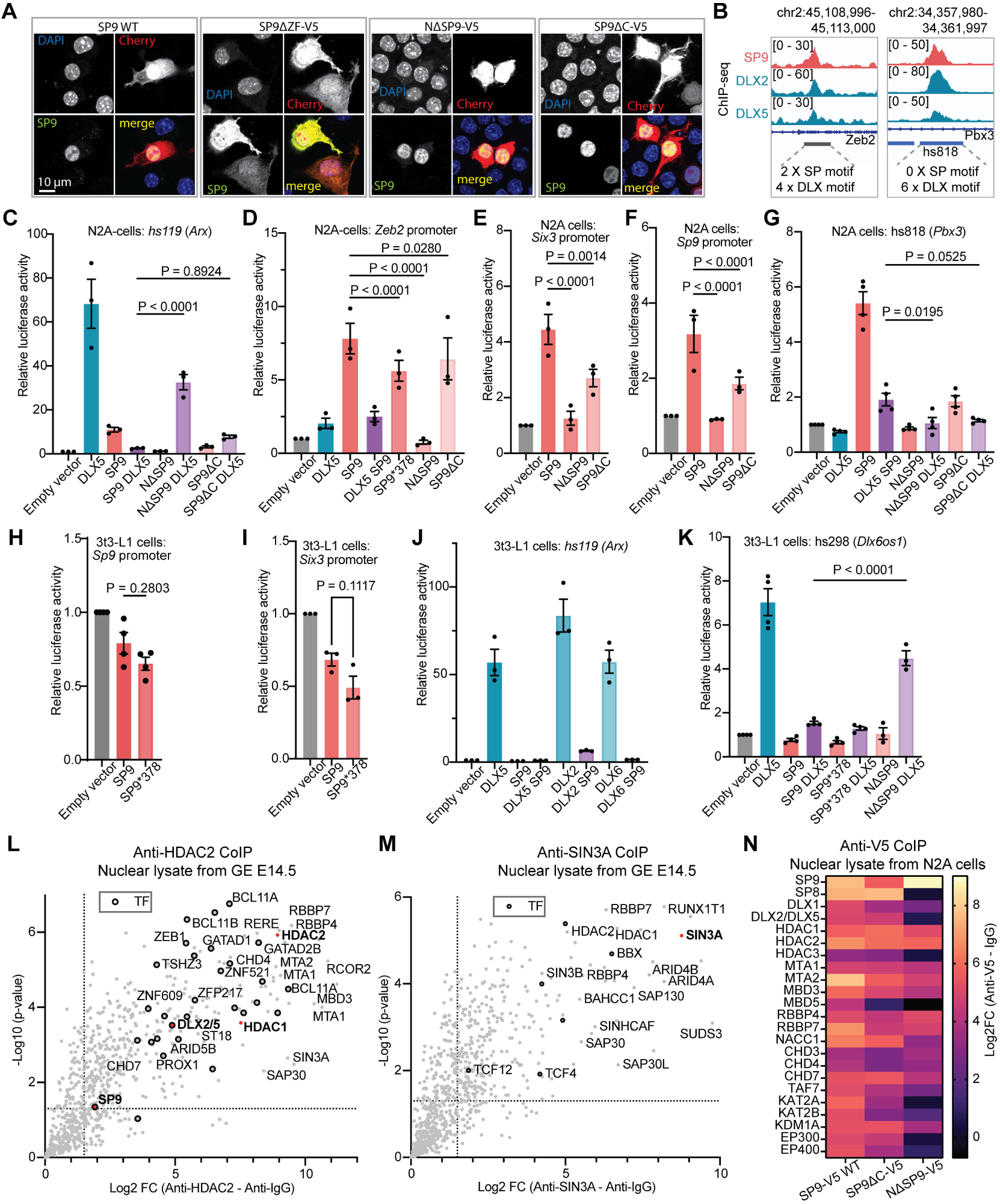
Interaction between SP9 and DLX5 at regulatory elements of GABAergic lineage genes, related to Figure 5. **A**, Representative confocal images of N2A cells co-transfected with mCherry (red) and WT or mutant SP9-V5 (green) constructs, immunostained with anti-V5 antibody. Nuclei were counterstained with DAPI (blue). Scale bar, 10 µm. **B**, Genome browser tracks showing ChIP-seq signal for SP9, DLX2, and DLX5 in E13.5 mouse GE at the *Zeb2* promoter (highlighted in grey) and the VISTA enhancer hs818 (highlighted in blue). Numbers below indicate the presence of SP and DLX motifs in the selected regulatory elements. Data from (Lindtner et al., 2019; Catta-Preta et al., 2025; Xu et al., 2018). **C-G**, Luciferase reporter activity in N2A cells, complementing the hs298 (*Dlx6os1*) data shown in Figure 5D, for the hs119 enhancer of *Arx* (**C**), *Zeb2* promoter (**D**), *Six3* promoter (**E**), *Sp9* promoter (**F**), and hs818 enhancer of *Pbx3* (**G**). Cells were co-transfected with combinations of *Dlx5*, *Sp9*, *N*Δ*Sp9*, *Sp9*378*, or *Sp9*Δ*C* as indicated. **H-K**, Luciferase reporter activity in 3T3-L1 cells for the *Sp9* promoter (**H**), *Six3* promoter (**I**), hs119 enhancer of *Arx* (**J**), and hs298 enhancer of *Dlx6os1* (**K**). Cells were co-transfected with combinations of *Dlx2*, *Dlx5*, *Dlx6*, *Sp9*, *Sp9*378*, or *N*Δ*Sp9* as indicated. **C-K**, Bars represent mean ± s.e.m. of 9 (C-F,I,J) or 12 (G,H,K) replicates from 3 (C-F,I,J) or 4 (G,H,K) independent batches performed in triplicate; points indicate batch means. Statistical significance was assessed by two-way ANOVA with Tukey’s honestly significant difference (HSD) post hoc test. Exact *P*-values are provided in Table S7. **L,M**, Volcano plots of CoIP-MS using anti-HDAC2 (**L**) or anti-SIN3A (**M**) on E14.5 mouse GE nuclear lysates, relative to IgG controls. Proteins with log_2_FC > 1.5 and *P* < 0.05 were considered significantly enriched (n = 375 in L, n = 290 in M). Transcription factors (TFs) are indicated in black. *P*-values were calculated using a two-sided Student’s t-test with permutation-based false discovery rate (FDR) correction. **N**, Heatmap of protein enrichment in anti-V5 CoIP-MS from N2A cells overexpressing SP9-V5 WT, SP9ΔC-V5, or NΔSP9-V5. Color indicates log_2_FC between anti-V5 and IgG control immunoprecipitates. Listed proteins include SP-family TFs (SP9, SP8), DLX factors (DLX1, DLX2/DLX5), NuRD complex components (HDAC1, HDAC2, HDAC3, MTA1, MTA2, MBD3, MBD5, RBBP4, RBBP7, NACC1, CHD3, CHD4, CHD7), transcriptional coactivators (KAT2A, KAT2B, KDM1A, EP300, EP400, TAF1).

**Figure S 7:**
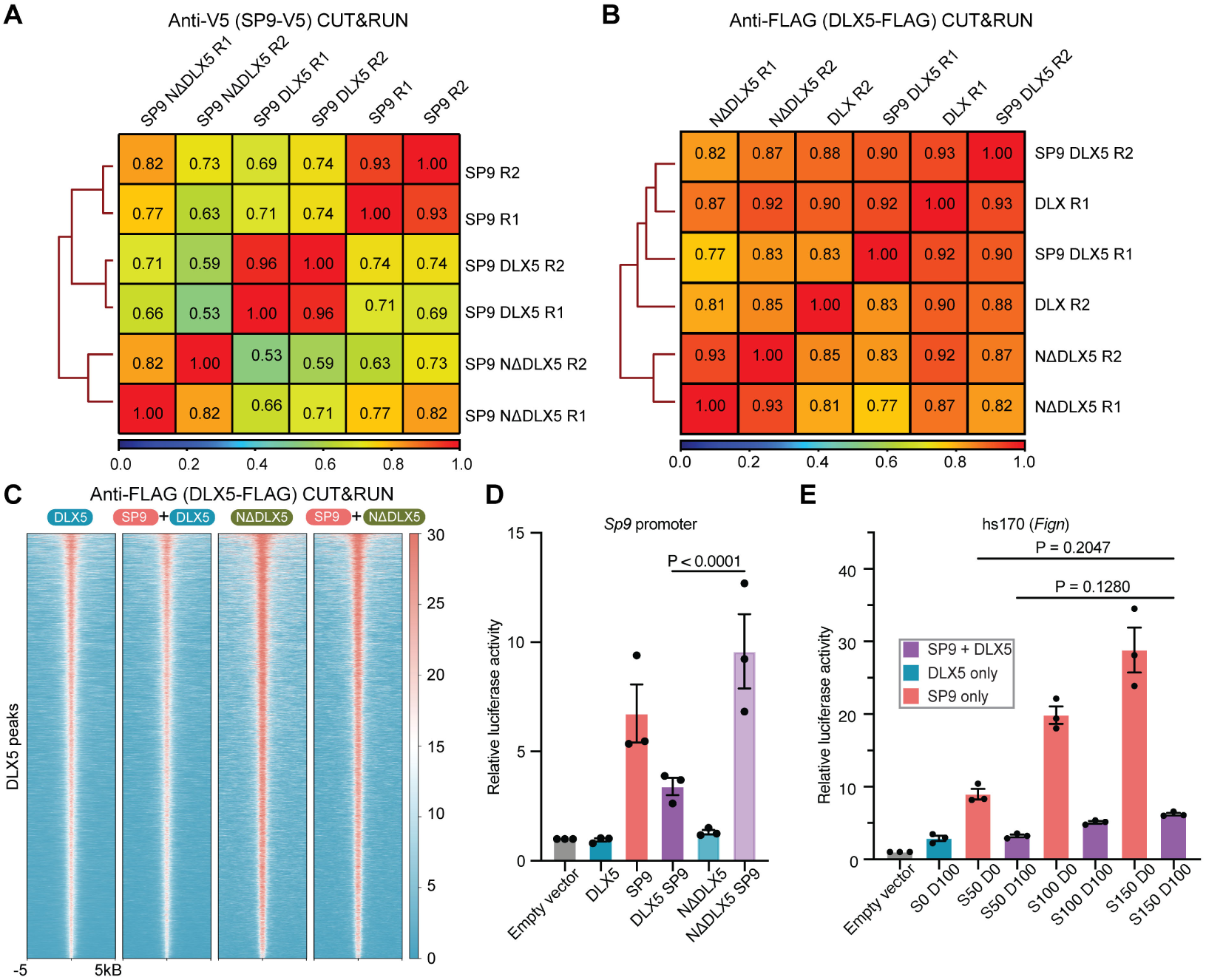
DLX5 controls SP9 genomic localization through its N-terminal domain, related to Figure 6. **A,B**, Genome-wide Spearman correlation matrices of CUT&RUN signal across overexpression conditions in N2A cells, based on 10 kb-binned BigWig signals. **A**: anti-V5 CUT&RUN profiling SP9-V5 occupancy. **B**: anti-FLAG CUT&RUN profiling DLX5-FLAG occupancy. Replicates cluster tightly, indicating high reproducibility. “R1” and “R2” denote biological replicates. **C**, Heatmaps of DLX5-FLAG CUT&RUN signal across DLX5-bound regions (±5 kb from peak center). Four conditions are shown: DLX5 (DLX5-FLAG) alone, SP9+DLX5 (SP9-V5+DLX5-FLAG), NΔDLX5 (NΔDLX5-FLAG) alone, and SP9+NΔDLX5 (SP9-V5+NΔDLX5-FLAG). **D**, Luciferase reporter activity driven by the *Sp9* promoter in N2A cells transfected with combinations of *Sp9*, *Dlx5*, or *N*Δ*Dlx5* as indicated. **E**, Luciferase reporter activity driven by the hs170 enhancer of *Fign* in N2A cells transfected with varying amounts (ng plasmid DNA) of *Sp9* (S) and *Dlx5* (D) to assess ratio-dependent transcriptional activation. **D,E**, Bars represent mean ± s.e.m. of 9 replicates from 3 independent batches performed in triplicate; points indicate batch means. Statistical significance was assessed by two-way ANOVA with Tukey’s honestly significant difference (HSD) post hoc test. Exact *P*-values are provided in Table S7.

## Methods

### Animals

All experiments were conducted according to institutional guidelines of the Max Planck Society and the regulations of the local government ethical committee (Beratende Ethikkommission nach §15 Tierschutzgesetz, Regierung von Oberbayern). All procedures were approved by the Bavarian government for the Max Planck Institute for Biological Intelligence (MPIBI) (ROB-55.2-2532.Vet_02-18-81 and ROB-55.2-2532.Vet_02-23-87). All mouse colonies were maintained in accordance with protocols approved by the Bavarian government. Mice were group-housed in isolated ventilated cages (room temperature 22 ^◦^C ± 1 ^◦^C, relative humidity 55 % ± 5 %) under a 12 h/12 h dark/light cycle with *ad libitum* access to food and water. Mouse strains used were the following: wild-type C57BL/6NRj (in-house breeding) and Cas9–EGFP (B6.Gt(ROSA)26Sor^tm1.1(CAG-cas9*,-EGFP)Fezh/J^, JAX, 026179) (Platt et al., 2014). Embryos were staged in days post coitus, with E0.5 defined as 12:00 of the day a vaginal plug was detected after overnight mating.

### Cell line

Mouse N2A neuroblastoma cells (Neuro2A cells, ECACC, 89121404), human HEK293FT (Thermo Fisher Scientific, R70007), mouse 3T3-L1 (CLS Cell line service, 400103) were cultured in Dulbecco’s modified Eagle medium (DMEM, Sigma, D6429) supplemented with 10% (v/v) fetal bovine serum (FBS, Sigma, F9665) and containing 1% (v/v) antibiotics (100 U ml^−1^ penicillin, 100 µg mL^−1^ streptomycin, Sigma, P0781). Cells were incubated at 37 ^◦^C in a 5% CO_2_ humidified atmosphere and passaged twice a week. Cell passage numbers were limited to no more than ten.

### tCROP-seq and TrackerSeq sample and library preparation

#### gRNA selection and vector construction

Single-guide RNAs (gRNAs) were designed using CRISPick for CRISPR knockout (Doench et al., 2016; Sanson et al., 2018) and evaluated with inDelphi (Shen et al., 2018) to maximize frameshift efficiency. Five gRNAs targeting *Sp9* were cloned into a piggyBac backbone using single-stranded DNA oligonucleotides (IDT) and NEBuilder HiFi DNA Assembly (NEB, E5520). The piggyBac vector (Addgene, 229995) encodes tdTomato and the gRNA under the human U6 promoter and contains a capture sequence in the gRNA scaffold for 10x Genomics feature barcode retrieval (cs1 incorporated at the 3’ end; (Replogle et al., 2020; Dvoretskova et al., 2024)). gRNA cutting efficiency was assessed in N2A cells. Cells were co-transfected with pCAG-Cas9-EGFP (gift from R. Platt) and gRNA plasmids using Lipofectamine 3000 (Thermo Fisher Scientific, L3000008). Forty-eight hours after transfection, tdTomato- and EGFP-positive cells were sorted using a Beckman Coulter CytoFLEX SRT. Genomic DNA was extracted using the Quick-DNA Miniprep Plus Kit (Zymo, D4068), and target regions were amplified using Q5 High-Fidelity DNA Polymerase (NEB, M0491S) with primers listed in Table S1. PCR products were subjected to Sanger sequencing (Eurofins), and knockout efficiency was quantified using TIDE (Brinkman et al., 2014). The knockout efficiency of the selected gRNA is summarized in Table S1. ZymoPURE II Plasmid Midiprep kit (Zymo Research, D4200) was used to produce endotoxin-free plasmid DNA for IUE.

#### TrackerSeq library preparation and validation

TrackerSeq is a piggyBac transposon-based (Ding et al., 2005) lineage tracing tool that is compatible with the 10x Genomics Chromium platform (Bandler et al., 2022). It records clonal lineages of single cells through the integration of oligonucleotide sequences into the genome of mitotic progenitors. Each lineage barcode is a 37 bp long synthetic nucleotide that consists of short random nucleotides bridged by fixed nucleotides. We followed the protocols from Bandler et al. to prepare TrackerSeq plasmids (Bandler et al., 2022). Briefly, an oligo library was cloned downstream of the Read2 partial primer sequence in the purified donor plasmid via Gibson Assembly reactions (NEB, E2611S). Gibson assembly reactions were then pooled and desalted with 0.025 *μ*m MCE membrane (Millipore, VSWP02500) for 40 min and concentrated using a SpeedVac (Eppendorf). A total of 3 *μ*l of the purified assembly was incubated with 50 *μ*l of NEB 10-*β*-competent *Escherichia coli* cells (NEB, C3019H) for 30 min at 4 ^◦^C, then electroporated at 1.95 kV, 200 Ω, 25 *μ*F (Bio-Rad, Gene Pulser Xcell Electroporation Systems).

Electroporated *E. coli* were recovered for 60 min shaking at 37 ^◦^C and then transferred to 3L LB, and grown in LB medium, supplemented with sucrose and ampicillin, up to optical density ≤ 0.5. The plasmid library was purified using 4 reactions of a column purification kit (Zymo Research, D4200). We first assessed the integrity of the TrackerSeq barcode library by sequencing it to a depth of 9.3 million reads to test whether any barcode was over-represented. After deduplication and error-aware clustering (allowing for a single error from the R2 FASTQ files) using Bartender (Zhao et al., 2018), 5.6 million unique lineage barcodes were recovered. Extrapolation of maximal library diversity based on the empirical diversity and representation at this sequence depth was performed using preseq, with an upper limit estimate of 12M barcodes. The read count distribution over barcodes was consistent with a 0-inflated Poisson distribution with very few outliers (i.e. barcodes with high read counts: the barcode with most reads had 58 reads, then 30), indicating uniform representation of barcodes.

#### Mice and in utero surgeries

Wild-type C57BL/6NRj females from in-house breeding were crossed with Cas9-EGFP males. Embryonic age was determined as days post coitum, with embryonic day 0.5 (E0.5) defined as noon on the day a vaginal plug was detected following overnight mating. Timed pregnant females were anesthetized with isoflurane (5% for induction and 2.5% during surgery, CP Pharma) and received Metamizol (WDT) for analgesia. *In utero* injections were performed using a Nanoject III programmable nanoliter injector (DRUM3-000-207). A total volume of 700 nl of DNA solution containing 0.6 *μ*g/*μ*l pEF1a-pBase (piggyBac transposase), 0.45 *μ*g/*μ*l Trackerseq barcode library and 0.8 *μ*g/*μ*l gRNA plasmid, diluted in sterile 0.9% NaCl with 0.002% Fast Green FCF (Sigma, F7252), was delivered into the lateral ventricle. The electroporation was carried out across the uterine wall by positioning the embryonic head between platinum-plated tweezer electrodes (5 mm diameter; BTX, 45-0489), followed by delivery of five square-wave pulses (35 V, 50 ms duration, 1 Hz) using an ECM830 electroporator (BTX) (Saito, 2006). Electrodes of this size were used to achieve broad targeting of the GE, including medial, lateral, and caudal regions. Prior to tissue processing for scRNA-seq, brains were inspected under a stereomicroscope. Only samples meeting the following criteria were included for downstream analysis: (i) widespread tdTomato-positive neurons in the neocortex, (ii) dense tdTomato labeling throughout the striatum, and (iii) tdTomato-positive neurons in the OB.

#### Sample collection and sequencing

Electroporated brains were collected from mouse embryos at E16.5 and maintained in ice-cold Leibovitz’s L-15 medium (Thermo Fisher Scientific, 21083027) supplemented with 5% FBS (Sigma, F9665). The same medium was used throughout flow cytometry sorting. Single-cell suspensions were generated using the Papain Dissociation System (Worthington, LK003150) according to a previously described protocol (Dvoretskova et al., 2024), with dissociation performed on a gentleMACS Octo Dissociator (Miltenyi Biotec). Immediately prior to sorting, DAPI (Thermo Fisher Scientific, D1306) was added to the cell suspension at a final concentration of 100 ng/mL to exclude dead cells. Fluorescence-activated cell sorting was performed on a Beckman Coulter CytoFLEX SRT using a 100 *μ*m nozzle to isolate tdTomato- and EGFP-positive, DAPI-negative cells. Following sorting, cells were centrifuged and resuspended in PBS (Corning, 21040) containing 0.02 % BSA (Gibco, 15260037) and immediately loaded onto the 10x Genomics Chromium platform for Gel Beads-in-Emulsion (GEM) generation and cDNA generation with cell- and transcript-specific barcodes. Single-cell 3’ gene expression and gRNA libraries were prepared using the Chromium Single Cell 3’ Reagent Kit v3.1 with Feature Barcoding technology (PN-1000268), following the manufacturer’s protocol (10x Genomics, CG000316). Libraries were constructed using the Chromium Single Cell 3’ Library Construction Kit v3.1 (PN-1000190) and the 3’ Feature Barcode Kit (PN-1000262) according to the manufacturer’s instructions. The TrackerSeq lineage libraries were amplified from 10X Genomics cDNA libraries with the Q5 polymerase (NEB, M094S) in a 50 *μ*l reaction, using 10 *μ*l of cDNA as template following Bandler et al. protocol (Bandler et al., 2022). Libraries were purified with a dual-sided selection using SPRIselect (Beckman Coulter, B23318). The quantification of the libraries was performed with the 4200 TapeStation. The resulting libraries were circularized, quantified, pooled, and sequenced on the AVITI System platform (Element Biosciences) at the Max Planck Institute of Biochemistry (MPIB) NGS Core Facility.

### tCROP-seq and scRNA-seq preprocessing

Raw FASTQ files for each replicate were concatenated and preprocessed using Cell Ranger count and aggr (10x Genomics, v8.0.1) with default parameters and user-provided capture sequence mapping for experimental gRNAs. The gLacZ data taken from (Dvoretskova et al., 2024) were also re-processed from raw FASTQ files using the same pipeline. Individual replicate datasets were preprocessed using scanpy [v1.12] (Wolf et al., 2018) according to the standard tutorial protocol with top features = 3000 and n_pcs = 50, and Scrublet was used to detect doublets (Wolock et al., 2019). Doublet cell barcodes were exported for removal from TrackerSeq lineage matrices and then removed from transcriptome single-cell files (see TrackerSeq methods). Cells with fewer than 500 gene features were removed, and genes with fewer than 3 non-zero count cells were removed. Clonal assignments from TrackerSeq analysis on each 10x sample were mapped into metadata prior to merging of replicates into an aggregate single-cell file. For replicate 1, since both gSp9 and gLacZ embryos were pooled on the same 10x reaction, we assigned cells to gLacZ or gSp9 only if RNA counts for gRNAs unambiguously indicated membership in only one guide condition. Cells with both guides detected were removed from downstream analysis and their cell barcodes exported for removal from the TrackerSeq lineage matrix, since these are likely produced by doublets.

All replicate adata files were merged in scanpy (Wolf et al., 2018), and doublets with a score ≥ 0.05 were removed. Raw counts were normalized and scaled. The default scanpy processing was performed again with top features = 3000 and n_pcs = 50, then knn and UMAPs were computed using default parameters. Since batch effects were present in the default UMAPs, we used harmony [v0.0.10] for batch correction of principal components (PCs), with the pooled replicate 1 serving as an internal control for batch vs. perturbation driven effects (Korsunsky et al., 2019). Leiden clustering performed on the resulting harmony UMAP and visualized on the original UMAP confirmed that cell groups exhibiting batch effects share the same transcriptomic phenotype in both UMAPs. Using Leiden clusters at a resolution of 1.0, marker gene sets were generated using rank_genes_groups with Wilcoxon rank sum tests and cells were manually annotated using marker genes annotated in mousebrain.org adolescent reference and previously published C. Mayer laboratory datasets (Zeisel et al., 2018; Mayer et al., 2018; Bandler et al., 2022; Dvoretskova et al., 2024) (Table 2). Clusters 20, 23, 24, 25, 26, and 27 exhibited markers for either immune or vascular cell types were removed from the dataset (Figure S1E). Cell cycle inference was performed using score_genes_cell_cycle based on mouse ortholog genes generated using gprofiler2 [v0.2.3] based on cell cycle genes and inference algorithm from Seurat human cell cycle list (Butler et al., 2018) (Figure S1F). Leiden clusters were re-computed on the neural cell datasets at resolutions from 0.5 to 2.0 and marker genes were generated for each set of Leiden clusters using rank_genes_groups with Wilcoxon rank sum tests. Detailed cell group annotations were performed for the full neural dataset (glutamatergic and GABAergic neurons) as well as the GABAergic only subset, using the same reference material (Figures 1B,C, S2A,C; Table 2).

Since *Sp9* is not expressed in the excitatory lineages, a GABAergic only dataset was generated by sub-setting all mitotic clusters and only post-mitotic clusters with both *Gad1* and *Gad2* expression (Figures 1E, S2A,C). Similar approaches were taken for further subsets of the GABAergic dataset, such as the PN or IN subsets which contain only terminally differentiated cell states and their immediate immature precursor states.

### MILO differential abundance analysis

Differential abundance (DA) testing was performed with MILO implemented via the pertpy package [v1.0.3] (Dann et al., 2022). The full GABAergic subset, PN and IN subsets, were used for MILO analysis using parameters as shown in Table S3. FDR thresholds and log2FC DA of individual neighborhoods were adjusted for each analysis, with cutoffs shown in the Table S3. These neighborhoods were selected for DE testing. DE was computed between enriched vs. depleted neighborhoods meeting the stated log2FC and FDR thresholds by pseudobulking gene expression by neighborhood (the sum of all counts from cells in the neighborhood) and performing pyDESeq2 [v0.5.2] (Muzellec et al., 2023) on neighborhoods as metacells. To assess the rate of potential false positives introduced by use of metacells with overlapping membership we also performed sample-pseudobulked DE between cells that were members of only positive vs. negative neighborhoods, and observed concordance for all genes but *Lingo2* and *Fam19a2* (Table 3). MILO of the clone2vec embedding space was performed similarly, but without a DE analysis of DA neighborhoods.

### PyDESeq2 differential expression analysis

Differential expression (DE) analysis was performed comparing gSp9 to gLacZ cells using pseudobulked counts by experimental replicate-gRNA with 2 replicates for the gSp9 condition and 3 replicates for the gLacZ control condition. The scRNA-seq data subsets for each DE analysis are shown in Table S4. Expression matrix columns for captured gRNAs and sex-linked genes were removed since sample inputs for each condition were not balanced by sex. Raw counts for each replicate-gRNA condition were aggregated as a sum using decoupler-py[v2.0.6] and replicates were assigned. PyDESeq2 was used to perform DE testing, and resulting *P*-values were corrected using Benjamini-Hochberg FDR (Muzellec et al., 2023). Volcano plots highlighted genes according to the log2FC and adj_p_value cutoffs shown in Table S4.

### Trajectory and pseudotime analysis

Pseudotime was computed on the GABAergic data subset using a diffusion pseudotime implementation by sc.tl.dpt using inh_IP_Abracl cluster as the root node (Haghverdi et al., 2016). We used PAGA to infer trajectories between cell states using cell connectivities computed from harmony principal component analysis (PCA) (Wolf et al., 2019). Based on the resulting trajectories, we oriented annotated cell states on the shared MSN trajectory spanning from annotated clusters inh_IP_Abracl to inh_PN_imm_MSN_Pou3f1, and assigned terminally differentiated D1 and D2 MSN clusters to distinct lineages. Cells contained within clusters that were categorized within these shared and distinct lineage trajectories were ordered by their pseudotime to plot gene expression on a continuous axis.

For plotting aggregate gene expression of D1 and D2 MSNs differentiation trajectories, we performed 1D Gaussian kernel to scaled and normalized counts over a 200 cells window, using ‘mirror’ method at data boundaries. Since the max pseudotime values for D1 MSNs were greater than D2 MSNs, likely due to arbitrary graph distance rather than meaningful real time disparities, we scaled the distinct pseudotime scores proportionally between the highest shared pseudotime and the highest D2 MSN pseudotime so that pseudotime values in the distinct lineage trajectories were relatively comparable and ending at 1.0.

### TrackerSeq preprocessing

Raw barcode library sequencing fastqs were preprocessed, merged, filtered, and assigned to clonal identities using the updated TrackerSeq pipeline available on the mayerlab github. In brief, reads are paired and filtered for P5, P7 index and 10X cellbarcode quality, cell barcodes are whitelisted using UMI_Tools whitelist and extract. Unique combinations of cell barcodes, lineage barcodes (LBCs), and read UMIs are extracted and quantified. Unique lineage barcodes and putative error reads are clustered using a pcr-aware algorithm with Bartender v1.1 (Zhao et al., 2018), with high quality clusters (4:1 ratio of centroid:error reads) used to map lineage barcode PCR/NGS errors to barcode centroids if their hamming distances were ≤ 5. LBCs detected for each cell were filtered for minimum read counts and minimum UMI counts of 10, and 5 respectively, and LBCs detected in only one cell were removed from the lineage matrix. An igraph network was constructed with edges between pairs of cells sharing at least one barcode; edges were weighted by the Jaccard similarity of LBC signatures between the two connected cells, where a score of 1 indicates that all detected lineage barcodes were present in both cells. Barcode collisions and sequencing errors typically yield low Jaccard scores, because only a small subset of LBCs is shared by chance. To assess robustness of the clonal assignments, we performed a parameter sweep, removing edges below thresholds ranging from 0.00 to 1.00 and comparing the resulting clone size distributions. These distributions remained stable down to thresholds as low as 0.2, indicating that the majority of clonal signatures were not driven by spurious barcode matches or overrepresented barcodes. For confidence, we used a Jaccard cutoff of >0.5 for the final clonal assignments. Connected components in the filtered graph were exported as clonal assignments. In total, 5,485 multicell clones were successfully matched into the scRNA-seq data, while 30,294 cells were assigned as single-cell clones.

### Clonal analysis

Clonal assignments were embedded into a latent space using the clone2vec algorithm (Erickson et al., 2025). All multi-cell clones were used in the embedding, single-cells with lineage barcodes were omitted. Since our dataset contained thousands of multicell clones, clonal context neighborhood size was set at 50. Training was performed over 100 epochs on 80% of clonal data with 20% withheld for validation, with stabilization of clone2vec vector loss after ∼30 epochs. Nearest neighbor connectivities were computed using knn of 15 and UMAP with standard parameters was used for 2D visualization of the clonal embedding. Clonal gRNA and replicate assignments were transferred from member cells. We removed 18 clones derived from experimental replicate 3 that contained pooled cells from both gLacZ and gSp9 embryos since clonal members contained gRNAs for both conditions, indicating false clonality likely driven by a multi-cell doublet from 10X barcoding. Additionally, 80 clones without sufficient gRNA counts in member cells were also removed, since lack of gRNA assignment could be driven by scRNA-seq dropouts rather than true absence of a guide. The resulting clonal embedding contains 5387 multi-cell clones ranging from 2 to 11 cells.

Unbiased clustering-based analysis: A Leiden clustering parameter sweep was performed with resolutions ranging from 0.5 to 2.5. We selected a clustering resolution of 1.5 for clustering analysis since this resolution provided the optimal separation of clonal output patterns across the continuously heterogeneous potency axes producing different terminal IN and PN states. After clustering, we quantified the proportion of gSp9 vs. gLacZ clones in each cluster.

Frequencies of cell type production were aggregated by clonal cluster (Figure S4B), and frequencies of clonal clusters were aggregated for each cell type to compare distributions of cell fates against the clustering of embedded clones (Figure 3G). To test whether PN fates shifted, we restricted analysis to clones producing at least one cell from the inh_PN_MSN_D1, inh_PN_MSN_D2, or inh_PN_ITC clusters and compared the per-clone cell-type frequency distributions between conditions using a Brunner-Munzel test on the pooled clone-level distributions (Figure 3J).

We aggregated clonal gene expression by summing the counts for all member cells. For genes of interest, clones with aggregate gene expression in the top or bottom 15th percentiles were categorized as “high” or “low” expressing clones, as shown in Figure S4C, and KDEs were generated on the transcriptomic embedding from member cells of these “high” and “low” categorized clones, shown in Figure S4D with bws ranging from 0.1 to 0.2.

### CatBoost gradient descent modeling to predict clone2vec embedding and clone cell-type production

Models were produced using aggregate clonal gene expression from either gLacZ or gSp9 clones as inputs: one set of models predicts clone2vec embedding vectors while the second set predicts cell type proportions in clones. All clone types were used as inputs. Training was performed using the CatBoost function from the clone2vec package with 20 percent of clones withheld from the training set and cell type annotations from the transcriptional embedding provided as categorical values to predict. This function performs gradient boosting to tune the model feature weights using the prediction loss at each training iteration. The trained models were used to generate SHAP values (SHapley Additive exPlanations) as explanatory feature weights for expressed genes that support model predictions of clonal clone2vec coordinates or cell types produced by a clone. SHAP values were produced for global models as well as for individual cell type predictions. In the case of specific cell type predictions, a correlation between gene features and cell types was also computed (Figure S5A). We contrasted the gLacZ and gSp9 based models by subtracting the SHAP values produced by the control gLacZ model from the gSp9 model values and plotting these changes in SHAPs against the gene’s correlation to cell fates of interest. Genes whose predictive weight decreased in the gSp9-trained model (ΔSHAP < 0) had higher predictive power for cell state in the presence of SP9, consistent with regulatory activity that depends on or operates in coordination with SP9 (Figures 3K, S5B). Conversely, genes with increased predictive weight in the gSp9-trained model (ΔSHAP > 0) became more informative in the absence of SP9, consistent with compensatory or SP9-independent regulation. Global SHAPs for the clone2vec vector prediction model were also contrasted between perturbation conditions. Since counts for the translation-deficient *Sp9* in the perturbed condition were similar compared to the gLacZ condition, its change served as an internal control for the robustness of comparison between perturbation conditions. It was reduced in both clone2vec and cell type global SHAPs.

### Luciferase assay

Regulatory elements were amplified from mouse genomic DNA using Q5 High-Fidelity DNA Polymerase (NEB, M0491) and primers listed in Table S6, and cloned into pGL4.24[luc2P/minP] (Promega, E8421) using the NEBuilder HiFi DNA Assembly Kit (NEB, E2621). *Dlx5* and *Sp9* coding sequences were amplified from mouse cDNA and cloned into pcDNA3.1 (GenScript). A C-terminal V5 tag was added to *Sp9* using NEBuilder HiFi DNA Assembly and a gBlock (IDT). C-terminal FLAG-tagged mouse *Dlx2*, *Dlx5*, and *Dlx6* expression vectors were obtained from GenScript. Mutant *Sp9* and *Dlx5* constructs were generated by PCR amplification followed by assembly using NEBuilder HiFi DNA Assembly with single-stranded DNA oligonucleotides (IDT). In the SP9 N-terminal domain mutant (NΔSP9), amino acids 2–186 were deleted; in the SP9 C-terminal domain mutant (SP9ΔC), amino acids 414–483 were deleted; and in the SP9 ZF domain mutant (SP9ΔZF), amino acids 333–414 were deleted. Luciferase reporter vectors were co-transfected with pNL1.1.PGK[Nluc/PGK] (Promega, N1441) as a normalization control in N2A cells or pNL1.1.TK[Nluc/TK] (Promega, N1501) in 3T3-L1 cells, together with different combinations of pcDNA3 (empty vector), pcDNA3-Dlx5, and pcDNA3-Sp9. N2A and 3T3-L1 cells were seeded in 48-well or 24-well plates at 50,000 or 80,000 cells per well, respectively, and transfected the following day using FuGENE 6 (Promega, E2691). Cells were harvested 24 h after transfection, and luciferase activity was measured using the Nano-Glo Dual-Luciferase Reporter Assay System (Promega, N1630) on a CLARIOstar microplate reader (BMG Labtech). NanoLuc activity was used for normalization. Statistical analyses were performed in GraphPad Prism (v11.0.0) using two-way ANOVA followed by Tukey’s honestly significant difference (HSD) test. Data were assumed to be normally distributed but were not formally tested. Statistical results are provided in Table S7.

### Immunocytochemistry in N2A cells

N2A cells were plated on poly-L-lysine (PLL; Sigma-Aldrich, P2636)-coated glass-bottom 24-well plates (Cellvis, P24-1.5H-N) at 50,000 cells per well and transfected the following day using FuGENE 6 (Promega, E2691) with an mCherry expression plasmid (Addgene, 41583) together with vectors expressing WT or mutant forms of SP9. Twenty-four hours post-transfection, cells were fixed in 4% PFA (Electron Microscopy Sciences, 15710-S) for 15 min at room temperature, permeabilized with 0.1 % Triton X-100 in PBS for 5 min, and blocked in 1 % goat serum (Sigma-Aldrich, G9023) and 2 % BSA (Sigma-Aldrich, A7030) in PBS for 1 h at room temperature. Cells were incubated with mouse anti-V5 primary antibody (Thermo Fisher Scientific, R96025; 1:400) diluted in blocking buffer for 1 h at room temperature, followed by three washes with PBS and incubation with goat anti-mouse Alexa Fluor 488 secondary antibody (Thermo Fisher Scientific, A11001; 1:500) for 1 h at room temperature in the dark. Coverslips were mounted using Fluoromount-G with DAPI (Invitrogen, 00-4959-52) and imaged on a Leica STELLARIS 5 DMi8 confocal microscope with a 63× oil-immersion objective (NA 1.4).

### Co-immunoprecipitation followed by Western blot

HEK293FT cells were plated at approximately 70 % confluency in 10-cm dishes. The following day, cells were transfected using TurboFect (Thermo Fisher Scientific, R0531) with expression constructs encoding *Dlx5*-FLAG and *Sp9*-V5, or *Sp9* variants (*N*Δ*SP9*-V5, *Sp9*Δ*ZF*-V5, *Sp9*Δ*C*-V5 and Sp9*378-V5). Illustrations of SP9 and DLX5 proteins were created using the Protein Domain Designer tool. Forty-eight hours post-transfection, cells were harvested. Cells were washed with PBS, collected by scraping, and pelleted by centrifugation at 400 × g for 5 min at 4 ^◦^C. Nuclear extraction was performed using ice-cold Buffer A (10 mM HEPES, pH 7.9; 10 mM KCl; 10 mM EDTA; 0.5 % Igepal; 1 mM DTT; Sigma, 20-265). Cells were incubated on ice for 10 min, briefly vortexed, and centrifuged at 800 x g for 10 min at 4 ^◦^C. The resulting nuclear pellet was resuspended in ice-cold PLB buffer (20 mM Tris-HCl, pH 7.5; 150 mM NaCl; 1.5 mM MgCl2; 1 mM DTT; 5 % glycerol; 1 % Triton X-100) supplemented with protease inhibitors (Roche, 11836170001) and phosphatase inhibitors (Roche, 4906845001). Because deletion of the ZF domain abolished nuclear localization of SP9 (Figure S6A), whole-cell lysates were used for these experiments instead of nuclear extracts (Figure 5B-C). Nuclear or whole-cell lysates were subjected to five cycles of sonication (30 s on / 30 s off at the highest setting) using a focused-ultrasonicator (Covaris E220). Lysates were clarified by centrifugation at 13,000 rpm for 15 min at 4 ^◦^C, and the supernatant was transferred to fresh, pre-chilled protein low binding tubes. For CoIP, clarified nuclear lysates were incubated with Anti-FLAG M2 Magnetic beads (Millipore, M8823) at 4 ^◦^C overnight with end-over-end rotation. Beads were then washed 2 times with ice-cold PLB buffer and 3 times with NET buffer (20 mM Tris-HCl, pH 7.2; 150 mM NaCl; 5 mM EDTA; 0.1% Triton X-100) to reduce nonspecific binding. Following the final wash, bound protein complexes were eluted by incubation with 4x LDS Sample Buffer (Thermo Fisher Scientific, NP0007) supplemented with 10× NuPAGE Reducing Agent (Thermo Fisher Scientific, NP0009) and heated at 90 ^◦^C for 7 min. Eluted samples were subsequently separated on 4–12% Bis-Tris gels (Thermo Fisher Scientific, NP0321) using 1× MES SDS running buffer (Merck, MPMES). Proteins were transferred onto PVDF membranes using Power Blotter Select Transfer Stacks (Thermo Fisher Scientific, PB5210) and Power Blotter Transfer Buffer (Thermo Fisher Scientific, PB7300) on a Power Blotter Station (Thermo Fisher Scientific). Membranes were blocked for 1 h at room temperature in 5% non-fat dry milk (Roth, T145.1) prepared in 1× TBS-T (BioLegend, 426309), followed by overnight incubation at 4 ^◦^C with primary antibodies diluted in the same blocking solution. The following antibodies were used: rabbit anti-SP9 antibody 1:3000 (Atlas Antibodies, HPA062616), rabbit anti-DLX5 1:5000 (Atlas Antibodies, HPA005670), mouse anti-V5 (Thermo Fisher Scientific, R96025). After three washes in TBS-T, membranes were incubated for 1 h at room temperature with Clean-Blot IP Detection Reagent (HRP; Thermo Fisher Scientific, 21230). Protein signals were detected using SuperSignal West Dura Extended Duration Substrate (Thermo Fisher Scientific, 34075) and visualized on an iBright Imaging System (Thermo Fisher Scientific, FL1000). Uncropped blot images related to Figures 4 are in Data S1.

### DNA pull-down/DNA affinity purification followed by mass spectrometry

DNA pull-down assays were adapted from Grawe et al. (2020) and Grand et al. (2021) and performed in triplicate (Gräwe et al., 2020; Grand et al., 2021). DNA oligonucleotides 46 bp long (wild-type and scrambled at the TAATT motif) were synthesized by IDT with 5’ biotinylation on the forward strand. In the scrambled control oligonucleotide, the AA nucleotides within each TAATT motif were replaced with CG. Oligonucleotide sequences are provided in Table S8. Oligonucleotides were annealed in annealing buffer containing 10 mM Tris-HCl (pH 7.5–8.0), 50 mM NaCl, and 1 mM EDTA. All subsequent steps were performed on ice or at 4 ^◦^C unless otherwise indicated. For each reaction, 20 *μ*l of Dynabeads M-280 Streptavidin (Thermo Fisher Scientific, 11205D) were equilibrated by washing twice with 1 ml of DNA-binding buffer (DBB; 10 mM Tris-HCl pH 8.0, 1 M NaCl, 1 mM EDTA, 0.05% Igepal). Annealed DNA oligonucleotides (2 *μ*M) were incubated with the beads in a final volume of 250 *μ*l DBB per replicate for 1 h at 4 ^◦^C with rotation. Beads were subsequently washed twice with DBB and once with protein-binding buffer (PBB; 50 mM Tris-HCl pH 8.0, 150 mM NaCl, 1 mM TCEP, 0.25% Igepal), supplemented with protease inhibitors (Roche, 11836170001), and resuspended in 100 *μ*l PBB per sample. Nuclear lysates were prepared from GEs dissected from E14–E15 C57BL/6 mouse embryos, flash-frozen in 1.5 ml tubes without any liquid, and stored at −80 ^◦^C until use. GEs from three independent litters were pooled for each experiment. Flash-frozen tissue was resuspended in 1 ml ice-cold PBS, dissociated by gentle pipetting, and centrifuged at 400 x g for 5 min at 4 ^◦^C. Nuclei were extracted using ice-cold Buffer A (10 mM HEPES pH 7.9, 10 mM KCl, 10 mM EDTA, 0.5% Igepal, 1 mM DTT), supplemented with protease inhibitors, incubated on ice for 10 min, briefly vortexed, and centrifuged at 800 × g for 10 min at 4 ^◦^C. The resulting nuclei were resuspended in 100 *μ*l per sample of Buffer B (20 mM HEPES pH 7.9, 400 mM NaCl, 1 mM EDTA, 1 mM DTT, 1% glycerol), supplemented with protease inhibitors, and incubated with agitation at 4 ^◦^C for 2 h. Nuclear extracts were clarified by centrifugation at 13,000 rpm for 15 min at 4 ^◦^C, and the supernatant was used for DNA pull-down assays. For each pull-down, 100 *μ*l of nuclear extract was incubated with oligonucleotide-bound beads in a total volume of 600 *μ*l PBB for 2 h at 4 ^◦^C with rotation. Beads were then washed three times with 1 ml PBB and twice with 1 ml PBS. After removal of residual PBS, beads were snap-frozen and stored at −80 ^◦^C until mass spectrometry (MS) analysis.

### Co-immunoprecipitation followed by mass spectrometry

The GEs were dissected from E14.5 mouse embryos in ice-cold PBS. The cells were dissociated by gentle pipetting and pelleted by centrifugation at 400 x g for 5 min at 4 ^◦^C. Nuclear extraction was performed using an ice-cold A buffer, supplemented with protease inhibitors (Roche, 11836170001). The cells were then incubated on ice at 4 ^◦^C for 10 min. The suspension was briefly vortexed, then centrifuged at 800 x g for 10 min at 4 ^◦^C. The nuclear pellet obtained was subsequently resuspended in ice-cold PLB buffer, supplemented with protease inhibitors (Roche, 11836170001) and phosphatase inhibitors (Roche, 4906845001). The nuclei were then subjected to five cycles of sonication (30 s on / 30 s off at the highest setting) using a Covaris E220. Lysates were clarified by centrifugation at 13,000 rpm for 15 min at 4 ^◦^C, and the supernatant was transferred to fresh, pre-chilled tubes. Then, 2 *μ*g of either rabbit anti-SP9 antibody (Atlas Antibodies, HPA062616; Figure 5F), rabbit anti-HDAC2 antibody (Abcam, ab32117; Figure S6L), rabbit anti-SIN3A antibody (Abcam, ab3479; Figure S6M), or rabbit anti-IgG antibody (Cell Signaling, 3900S; as a negative control across all conditions) was added, and the tubes were rotated at 4 ^◦^C overnight. Then, a 25 *μ*l slurry of Protein A/G magnetic beads (Thermo Fisher Scientific, 88802) was washed in NET buffer (50 mM Tris-HCl pH 7.5, 150 mM NaCl, 5 mM EDTA, 0.1% Triton X-100), blocked in 2 % BSA-NET buffer for 10 min, and then sequentially resuspended in ice-cold PLB buffer. The beads were added to the nuclear lysates and incubated for an additional 2 h. The beads were washed twice in ice-cold PLB buffer, twice in NET buffer, and twice in PBS. Finally, all liquid was removed and the samples were sent to the MPIBI MS facility. Proteins in the MS experiments were considered to be significantly enriched in the eluates when *P*-value < 0.05 and FC > 1.5.

For CoIP-MS experiments in N2A cells, cells were plated at approximately 70% confluency in 10-cm dishes and transfected the following day using TurboFect (Thermo Fisher Scientific, R0531) with expression constructs encoding WT *Sp9*-V5, *N*Δ*Sp9*-V5 or *Sp9*Δ*C*-V5 (i, Figure S6N) or with expression constructs encoding *Sp9*-V5 alone or co-expressing *Dlx5*-FLAG and *Sp9*-V5 (ii, Figure 6I-J). Nuclear lysates were prepared as described above, with the exception that PLB buffer in (ii) was supplemented with Turbo Nuclease (Benzonase; Jena Bioscience, EN-180S; 1 µL/mL) and poly-L-glutamic acid (PGA; Sigma-Aldrich, P4761; 0.4 mg/mL) to degrade nucleic acids and reduce nonspecific protein interactions (Hall Hickman and Jenner, 2024). CoIP was performed using anti-V5 antibody (Thermo Fisher Scientific, R96025) or mouse IgG2a isotype control (Merck, MABF1080Z), followed by MS.

### Liquid Chromatography-Mass Spectrometry (LC-MS)

#### Sample preparation

The beads were incubated at 37 ^◦^C for 20 minutes with SDC buffer (100 *μ*L, containing 1% sodium deoxycholate, SDC, Sigma-Aldrich; 40 mM 2-chloroacetamide, CAA, Sigma-Aldrich; and 10 mM Tris(2-carboxyethyl)phosphine, TCEP, Thermo Fisher Scientific; in 100 mM Tris, pH 8.0). Next, the samples were diluted with 100 *μ*L of milliQ water, and the proteins were digested overnight at 37 ^◦^C by adding 0.5 *μ*g of trypsin (Promega). The resulting supernatant was collected using a magnetic rack and acidified with trifluoroacetic acid (TFA, Merck) to a final concentration of 1 %. Any precipitated SDC was removed via centrifugation, and peptides (ca. 200 ng) were directly loaded onto Orbitrap Exploris 480 (Thermo Scientific) or Evotips (Evotip Pure, Evosep).

#### LC-MS data acquisition

##### LC-MS data acquisition for DNA affinity purification followed by MS (Figure 5E)

The Orbitrap Exploris 480 (Thermo Scientific) was operated in data-dependent acquisition (DDA) mode. Full MS survey scans were acquired over an m/z range of 300–1650 at a resolution of 60,000 (at m/z 200). The top 10 most intense precursor ions were selected for fragmentation by higher-energy collisional dissociation (HCD) using a normalized collision energy of 28. MS/MS spectra were recorded at a resolution of 15,000 (at m/z 200). The automatic gain control (AGC) targets were set to 3 × 10^6^ for MS1 and 1 × 10^5^ for MS2 scans, with maximum injection times of 100 ms and 60 ms, respectively.

##### LC-MS data acquisition for CoIP-MS experiment in Figure 5F

Peptides were eluted from the Evotips onto a 15 cm PepSep C18 column (1.5 *μ*m, Bruker Daltonics) using the Evosep One HPLC system (Evosep). The column was maintained at 50 ^◦^C, and peptide separation was achieved using the 30 SPD method. Eluted peptides were directly ionized and introduced into a timsTOF Pro mass spectrometer (Bruker) via electrospray ionization. Data acquisition was performed in data-independent acquisition (DIA) PASEF mode via timsControl. MS covered a scan range of 100–1700 m/z, and ion mobility ranged from 1/K0 = 0.70 to 1.30 *Vs* × *cm*^2^. The dual TIMS analyzer utilized equal ion accumulation and ramp times of 100 ms each, with a spectra rate of 9.52 Hz. For DIA-PASEF scans, the mass scan range was 350.2–1199.9 Da, and ion mobility ranged from 1/K0 = 0.70 to 1.30 *Vs* × *cm*^2^. Collision energy was linearly ramped based on ion mobility, from 45 eV at 1/K0 = 1.30 *Vs* × *cm*^2^ to 27 eV at 1/K0 = 0.85 *Vs* × *cm*^2^. A total of 42 DIA-PASEF windows were acquired per TIMS scan, with switching precursor isolation windows, resulting in an estimated cycle time of 2.21 seconds.

##### LC-MS data acquisition for CoIP-MS experiments in Figures S6L-N, S7I-J)

LC–MS/MS analyses were performed using an EASY-nLC 1200 system coupled to an Orbitrap Exploris 480 mass spectrometer (Thermo Fisher Scientific). Peptides were separated on a 30 cm analytical column (75 *μ*m inner diameter) packed in-house with 1.9 *μ*m ReproSil-Pur C18-AQ resin (Dr. Maisch GmbH). The column was maintained at 60 ^◦^C. Samples were loaded in buffer A (0.1 % (v/v) formic acid in water) and eluted at a flow rate of 250 nL min^−1^. Buffer B was increased linearly from 5 % to 30 % over 40 min, followed by a ramp to 95 % over 10 min and a final hold at 95 % for 10 min.

#### Data analysis

##### Data analysis for DNA affinity purification followed by MS (Figure 5E)

Raw data were processed using the MaxQuant computational platform with standard settings applied. Shortly, the peak list was searched against the Uniprot database of mouse (55466 entries, downloaded in 2023). Cysteine carbamidomethylation was set as static modification, and methionine oxidation and N-terminal acetylation as variable modifications. The match-between-run option was enabled, and proteins were quantified across samples using the label-free quantification algorithm in MaxQuant generating label-free quantification (LFQ) intensities.

##### Data analysis for CoIP-MS experiments (Figures 5F, S6L-N, S7I-J)

Raw data were analyzed with Spectronaut (v19 or 20, Biognosys) in directDIA+ (library-free) mode. The data was searched against a predicted library of the mouse database from uniprot (downloaded in 2023) and additional sequences of interest. Cysteine carbamidomethylation was set as static modification, and methionine oxidation and N-terminal acetylation as variable modifications. The protease was set to Trypsin/P.

Protein group output files generated by MaxQuant or Spectronaut were loaded into Perseus (version 2.0.11.0). The matrix was filtered to remove all proteins that were potential contaminants, only identified by site and reverse sequences. The LFQ intensities were then transformed by log2(x), and individual intensity columns were grouped by experiment. Proteins were filtered to keep only those with a minimum of three valid values in at least one group. Missing values were randomly imputed with a width of 0.3 and downshift of 1.8 from the standard deviation. To identify significant differences between groups, two-sided Student’s t-tests were performed with a permutation-based FDR of 0.05. GO enrichment analysis was performed using Enrichr to query the Gene Ontology (GO Molecular Function 2025) (Xie et al., 2021).

### ChIP-seq Data processing and analysis

Publicly available ChIP-seq datasets for SP9 (Xu et al., 2018; Catta-Preta et al., 2025), DLX2 (Lindtner et al., 2019), GTF2I (Kopp et al., 2020), histone modifications (H3K4me3, H3K4me1, and H3K27ac) (Lindtner et al., 2019) and NuRD complex (MBD3, CHD4, RBBP4, RBBP7, HDAC1, and HDAC2) (Price et al., 2022) were used in this study. All ChIP–seq datasets were processed using the seq2science [v1.2.4] (van der Sande et al., 2023) pipeline with default parameters. All ChIP–seq datasets were mapped to reference genome mm10.

SP9 Peaks were annotated with genomic features and associated TF binding motifs using the annotatePeaks.pl function from HOMER [v5.1] (Heinz et al., 2010). A Custom motif file with SP and DLX motifs was used to specifically assess the genomic binding distribution of these factors.

To characterize the chromatin context and co-binding patterns at SP9-bound regions, signal intensities for SP9, DLX2, GTF2I, histone modifications (H3K4me3, H3K4me1, and H3K27ac) and NuRD complex (MBD3, CHD4, RBBP4, RBBP7, HDAC1, and HDAC2) were computed using the computeMatrix function from deepTools (Ramírez et al., 2016). Normalized ChIP-seq coverage files (RPKM) were used as input and signal was calculated in reference-point set to peak center. The resulting score matrix was visualized using plotHeatmap function from deepTools (Ramírez et al., 2016) with k-means clustering to generate heatmaps.

#### 0.0.1 D2-SPN marker gene promoters and motif occupancy analysis

D2 MSN genes used in figure 4G were obtained by intersecting wilcoxon-rank-sum derived differential expression for enriched vs. depleted MILO-neighborhoods (as metacells) with single-cell markers for inh_PN_MSN_D2 clusters. The top 100 genes ranked by absolute MILO differential expression value were selected for analysis. Promoter regions were defined as ±200 bp around TSSs using NCBI RefSeq (mm10) annotations. SP9 ChIP-seq signal across promoter regions was quantified using deepTools computeMatrix, and promoters were classified into high- and low-occupancy groups based on the median promoter signal. High-occupancy promoters were subjected to motif analysis using HOMER [v5.1] annotatePeaks and a custom motif library containing SP-family and DLX-family motifs.Promoters were subsequently categorized according to the presence of SP-family motifs, DLX-family motifs, or both, and motif class distributions were visualized as stacked bar plots.

### CUT&RUN in N2A cells

#### CUT&RUN sample and library preparation

For CUT&RUN experiments, N2A cells were plated at 400,000 cells per well in 6-well plates and transfected the following day using Lipofectamine 3000 (Thermo Fisher Scientific, R0531) with expression constructs encoding *Sp9*-V5 or *Dlx5*-FLAG alone, or co-expressing *Dlx5*-FLAG and *Sp9*-V5 (overexpression experiment 1, OE1). In a second overexpression experiment (OE2), cells were transfected with *Sp9*-V5 or *N*Δ*Dlx5*-FLAG alone, or co-expressing *N*Δ*Dlx5*-FLAG and *Sp9*-V5. After 24 hours, we performed CUT&RUN using the CUTANA CUT&RUN Kit (EpiCypher, 14-1048) according to the manufacturer’s instructions. Cells were harvested by washing with PBS and detached with Trypsin-EDTA (0.025 % in PBS; Thermo Fisher Scientific, 25200056). Trypsin was neutralized with 1 % BSA in PBS, and cells were collected by centrifugation at 400 x g for 5 min. For each sample, 500,000 cells were processed and incubated with the following antibodies: Anti-V5 (SP9) (Thermo Fisher Scientific, R96025), Anti-FLAG (DLX5) (EpiCypher, 13-2031), Anti-H3K4me3 (EpiCypher, 13-0041), and Anti-IgG (EpiCypher, 13-0042) as a negative control. Two biological replicates were included for each antibody condition. After antibody binding, chromatin digestion and release were performed using pA-MNase according to the standard CUT&RUN protocol. DNA fragments were purified and used for sequencing library preparation with the NEBNext Ultra II DNA Library Prep Kit for Illumina (NEB, E7645S). Libraries were prepared following the manufacturer’s protocol and amplified by PCR. The resulting libraries were quantified, pooled, and subjected to paired-end sequencing on the AVITI System platform (Element Biosciences).

#### CUT&RUN Data processing and analysis

Raw paired-end FASTQ files were first evaluated for sequencing quality using FastQC [v0.11.9] (Andrews, 2010). Data processing was done based on the CUT&Tag tutorial by Zheng et al. (Zheng et al., 2020). Reads were aligned to the reference genome mm10 using Bowtie2 [v2.4.2] (Langmead and Salzberg, 2012). Alignment files were converted and processed using SAMtools [v1.9] (Li et al., 2009) to generate sorted BAM files for downstream analyses. Library quality was assessed based on fragment size distribution and alignment statistics.

Enriched chromatin regions were identified using MACS2 [v2.2.8] (Zhang et al., 2008) by keeping IgG as control. Peaks were called using a *P*-value threshold of 1 × 10^−4^ (-p 1e-4). To obtain high-confidence and reproducible peak sets, Irreproducible Discovery Rate (IDR) analysis [v2.0.3] was performed on biological replicates using the IDR framework. Peaks from replicate experiments were ranked by signal intensity (signal.value), and reproducible peaks were identified using an IDR threshold of 0.08. Resulting IDR-filtered peak sets from different conditions were merged using BEDTools [v2.30.0] to generate a unified peak set. Genome-wide coverage tracks were generated from BAM files using the bamCoverage function from deepTools [v3.5.0] (Ramírez et al., 2016) to produce RPKM-normalized bigWig files. These signal tracks were used for downstream visualization and comparative analysis across samples. For heatmap generation (Figure 6C), IDR-filtered SP9 peaks from the SP9+ and SP9+DLX5+ conditions were merged to create a unified peak set. Signal intensities were computed using computeMatrix function from deepTools, followed by k-means clustering using plotHeatmap. The clustered BED file generated for Figure 6C was used as input for downstream analysis. SP9 binding signal across the different conditions was computed using computeMatrix, and coverage plots were generated using the plotProfile function. Motif enrichment analysis was carried out using HOMER [v5.1] findMotifsGenome function (Heinz et al., 2010) to identify sequence motifs overrepresented within the clustered regions. Genomic feature annotation was then performed using annotatePeaks. Genome browser track profiles were generated using fluff profile from fluff [v3.0.4] (Georgiou and van Heeringen, 2016).

### Published data used in this study

GSE124936 (H3K27ac, H3K4me1, H3K4me3, DLX2 and DLX5 ChIP-seq from E13.5 GE (Lindtner et al., 2019)), GSE222183 (SP9 ChIP-seq from E13.5 GE (Xu et al., 2018; Catta-Preta et al., 2025)), GSE194076 (MBD3, CHD4, RBBP4, RBBP7, HDAC1 and HDAC2 ChIP-seq from E13.5 GE (Price et al., 2022)), GSE138234 (GTF2I ChIP-seq from E13.5 brain (Kopp et al., 2020)), and GSE231779 (gLacZ-condition TrackerSeq and tCROP-seq from E16.5 (Dvoretskova et al., 2024)) were downloaded from https://www.ncbi.nlm.nih.gov/geo. VISTA enhancer images were downloaded from the VISTA Enhancer Browser (https://enhancer.lbl.gov) (Visel et al., 2007). Genome browser track profiles in Figures 4I and S6B were generated using Integrative Genomics Viewer (IGV_2.19.7, (Robinson et al., 2011). Protein domain architectures were designed using the Protein Domain Designer tool (https://domaindesigner.farnunglab.com/).

## Resource availability

### Lead contact

Further information and requests for resources and reagents should be directed to and will be fulfilled by the lead contact, Christian Mayer.

### Data availability

Sequencing data generated in this study (single-cell RNA-seq and CUT&RUN) will be deposited in the Gene Expression Omnibus (GEO; CUT&RUN: GSE331084) and made publicly available upon publication. Mass spectrometry data have been deposited in MassIVE/ProteomeXchange under accession PXD078811 and will be made publicly available upon publication. Data are available from the lead contact upon reasonable request.

### Code availability

All custom code used for data processing, analysis, and figure generation will be made publicly available through a permanent repository with an archival DOI upon publication, and is available from the lead contact upon reasonable request.

## Acknowledgements

We thank members of the C. Mayer laboratory and R. Bandler (Yale University) for feedback and discussion; T. Lamprecht and A. Rawal (MPIBI) for technical support; J. Kuhl (somedonkey.com) for illustrations; R. H. Kim and members of the MPIB NGS Core Facility (RRID:SCR_025746); M. Spitaler and M. Oster from the MPIB Imaging Facility (RRID:SCR_025739); C. Polisseni and MPIBI Imaging Core Facility (RRID:SCR_026797); B. Steigenberger and members of the MPIB Mass Spectrometry Core Facility (RRID:SCR_025745); and members of the MPIB/MPIBI animal facility for their technical expertise. This work was supported by the Max Planck Society (Max Planck Research Group, Free-Floater Programme; to C.M.); the Deutsche Forschungsgemeinschaft (DFG, project ID 549328218; to C.M.); the Simons Foundation Autism Research Initiative (SFARI Pilot Award, SFI-AN-AR-Pilot-00009814; to C.M.); the Nancy Lurie Marks Family Foundation (Research Grant; to C.M.); and the European Molecular Biology Organization (EMBO Young Investigator Programme; to C.M.). I.A. and S.I. were supported by the ERC Synergy grant (KILL-OR-DIFFERENTIATE), Swedish Research Council, Paradifference Foundation, Bertil Hallsten Research Foundation, Cancer Foundation in Sweden, Knut and Alice Wallenberg Foundation, Austrian Science Fund. P.V.K. was supported by the ERC Synergy grant (KILL-OR-DIFFERENTIATE) and the Visiting Fellowship from the Austrian Academy of Sciences.

## Author contributions

Conceptualization, E.D., C.L., A.R.B., and C.M.; Methodology, E.D., C.L., A.R.B., S.I., P.K., and I.A.; Investigation, E.D., C.L., and A.R.B.; Formal analysis, E.D., C.L., A.R.B., and Y.K.B.; Resources, S.I., P.K., and I.A.; Supervision, I.A. and C.M.; Writing – original draft, E.D., C.L., A.R.B., and C.M.; Writing – review & editing, all authors; Funding acquisition, C.M.

## Declaration of generative AI and AI-assisted technologies in the writing process

During the preparation of this work, the authors used DeepL, Claude.ai, and ChatGPT in order to improve the language and readability of the manuscript. After using these tools, the authors reviewed and edited the content as needed and take full responsibility for the content of the publication.

## Declaration of interests

The authors declare no competing interests.

## Supplemental Material

Tables are presented as individual Excel files:

**Table 1:** sgRNA information and tCrop-seq/TrackerSeq experimental numbers.

**Table 2:** Marker gene lists for all scRNA-seq datasets and intersection of MILO DE with D2 MSN markers

**Table 3:** MILO neighborhood DE analysis for GABAergic, PN, and IN data subsets

**Table 4:** Results of differential expression analysis with pyDESeq2.

**Table 5:** ChIP-seq motifs enrichment analysis.

**Table 6:** Luciferase reporter vector information.

**Table 7:** Luciferase data and statistics.

**Table 8:** Oligonucleotides used in DNA-pull down experiment.

**Table 9:** Mass spectrometry analysis results from DNA pull-down and CoIP experiments.

**Data S1**: Uncropped blot images, related to Figures 4.

## Notes

### Competing Interest Statement

The authors have declared no competing interest.

### Summary of Updates

This version of the manuscript has been revised to update the title, add a new figure panel, and update several figures, with accompanying text edits to the abstract and main text for clarity. The underlying data, analyses, and conclusions are unchanged.

